# Structural Basis of Glycoform Selectivity in Prion Strains

**DOI:** 10.1101/2025.11.04.686259

**Authors:** Francesca Peccati

**Affiliations:** Center for Cooperative Research in Biosciences (CIC bioGUNE), Basque Research and Technology Alliance (BRTA) Bizkaia Technology Park, 48160 Derio, Spain; Ikerbasque, Basque Foundation for Science, 48013 Bilbao, Spain

## Abstract

Prion diseases originate from the pathological misfolding of the cellular sialoglycoprotein prion protein (PrP^C^), universally found across mammalian species, into an aberrant conformation termed PrP^Sc^, which exhibits high aggregation propensity and neurotoxicity. Distinct conformations of the misfolded and aggregated PrP^Sc^, termed prion strains, can cause different disease phenotypes and transmission characteristics. Different prion strains exhibit well-defined and distinct glycoform preferences arising from two sialylated, N-linked glycans. Glycosylation, and in particular sialylation, have been demonstrated to modulate the replication rate of PrP^Sc^, with profound implications for the propagation of prion diseases. In this work, we leverage high-resolution cryo-EM structural data and all-atom molecular dynamics simulations to elucidate the molecular basis of the glycoform preferences in mouse strains RML and ME7. We show that these preferences are determined by differential engagement of the major basic patch and palindromic region of PrP, shedding light on a long elusive, fundamental aspect of prion biology.

## Introduction

The central event underlying the transmission of prion diseases—such as bovine spongiform encephalopathy in cattle, chronic wasting disease in deer and elk, and Creutzfeldt–Jakob disease in humans—is the misfolding and aggregation of the host-encoded, GPI-anchored prion protein (PrP^C^) into *β*-rich fibrils, converting it into the abnormal PrP^Sc^ form that accumulates in the brain.^1,2^ Substantial evidence supports the *protein-only hypothesis*, which posits that PrP alone can self-replicate by progressively converting PrP^C^ through an autocatalytic templating mechanism.^3,4^ According to this model, interspecies transmissibility depends on the ability of an exogenous PrP^Sc^ seed to induce PrP^C^ conversion in the host species. A key characteristic of prion diseases—also thought to underlie other, more prevalent amyloid-related disorders such as Alzheimer’s disease^5^—is the strain phenomenon, namely that PrP can misfold into multiple aggregated conformations, giving rise to distinct disease phenotypes.^6–10^ From a molecular perspective, interspecies transmission therefore depends, to a first approximation, on amino-acid sequence compatibility between species—that is, on the ability of the PrP^C^ from a given species to adapt to the various aggregated PrP^Sc^ forms (strains) of a different species.^8^

However, extensive evidence indicates that, beyond sequence compatibility, post-translational modifications—particularly glycosylation—strongly influence prion disease transmission across species.^2^ PrP^C^ contains two conserved N-glycosylation sites (residues 180 and 196 in mouse numbering), giving rise to four glycoforms: diglycosylated, monoglycosylated at either site (sharing the same moleculat weight), and unglycosylated. While the ratios of these glycoforms remain stable in PrP^C^, they vary widely among PrP^Sc^ strains, suggesting a connection between glycosylation state and strain-specific infectivity.^2,11,12^ Anti-PrP Western blots of PrP^Sc^ typically display three bands corresponding to the di-, mono-, and unglycosylated forms. The pattern and intensity of these bands are strain-specific and serve to differentiate strains, indicating that glycoform selectivity is encoded in the structural distinctions among them.^13^ A strong correlation also exists between glycan composition—which is in-trinsically heterogeneous—and the distribution of glycoforms, with a clear anticorrelation between glycan sialylation level and the abundance of the diglycosylated form.^13^ Sialic acid (N-acetylneuraminic acid, Sia) carries a negative charge at neutral pH; thus, the exclusion of diglycosylated, hypersialylated PrP^C^ units from the growing PrP^Sc^ fibril has been attributed to electrostatic repulsion between charges of the same sign,^14^ potentially modulating both the thermodynamic and kinetic factors that govern prion propagation.

Although these phenomena are well established, a molecular interpretation of strain differences and the role of glycosylation has long remained elusive, primarily due to the inherent difficulty of determining PrP^Sc^ structures at high resolution. The recent determination of quasi-atomic resolution structures of full-length, ex vivo PrP^Sc^ from distinct strains by cryo-EM (hamster 263K,^15^ mouse RML,^16,17^ mouse ME7,^18^ mouse 22L,^19^ and white-tailed deer CWD^20^) represents a major advance in the structural understanding of prion diseases. The present work focuses on the mouse strains RML and ME7 (Figure 1A). These strains exhibit broadly similar architectures, characterized by conformationally conserved N-terminal and C-terminal lobes forming a surface groove connected by a conformationally variable segment spanning residues 112–130.^18^ This flexible region differs markedly between the two strains, producing a large variation in groove width—wider in RML and narrower in ME7.^18^ The N180 glycosylation site lies at the bottom of the groove, whereas the N196 site lies at the tip of the C-terminal lobe.^18^ The 100–110 segment, rich in lysine residues, forms the major basic patch of PrP^Sc^ on the N-terminal side of the groove;^18^ this region carries a high positive charge density (see electrostatic potential maps in Figure 1A) and is decorated by phosphotungstate (PTA) polyanions in the cryo-EM structures, consistent with a strong tendency to interact with negatively charged species. The glycan at N180 also lies near the AGAAAAGA palindrome (residues 112–119) within the conformationally variable region (Figure 1B). This segment is essential for both the formation of the PrP^Sc^ structure and the PrP^Sc^/PrP^C^ interaction required for fibril propagation, as its deletion generates a dominant-negative inhibitor.^21^

**Figure 1.**
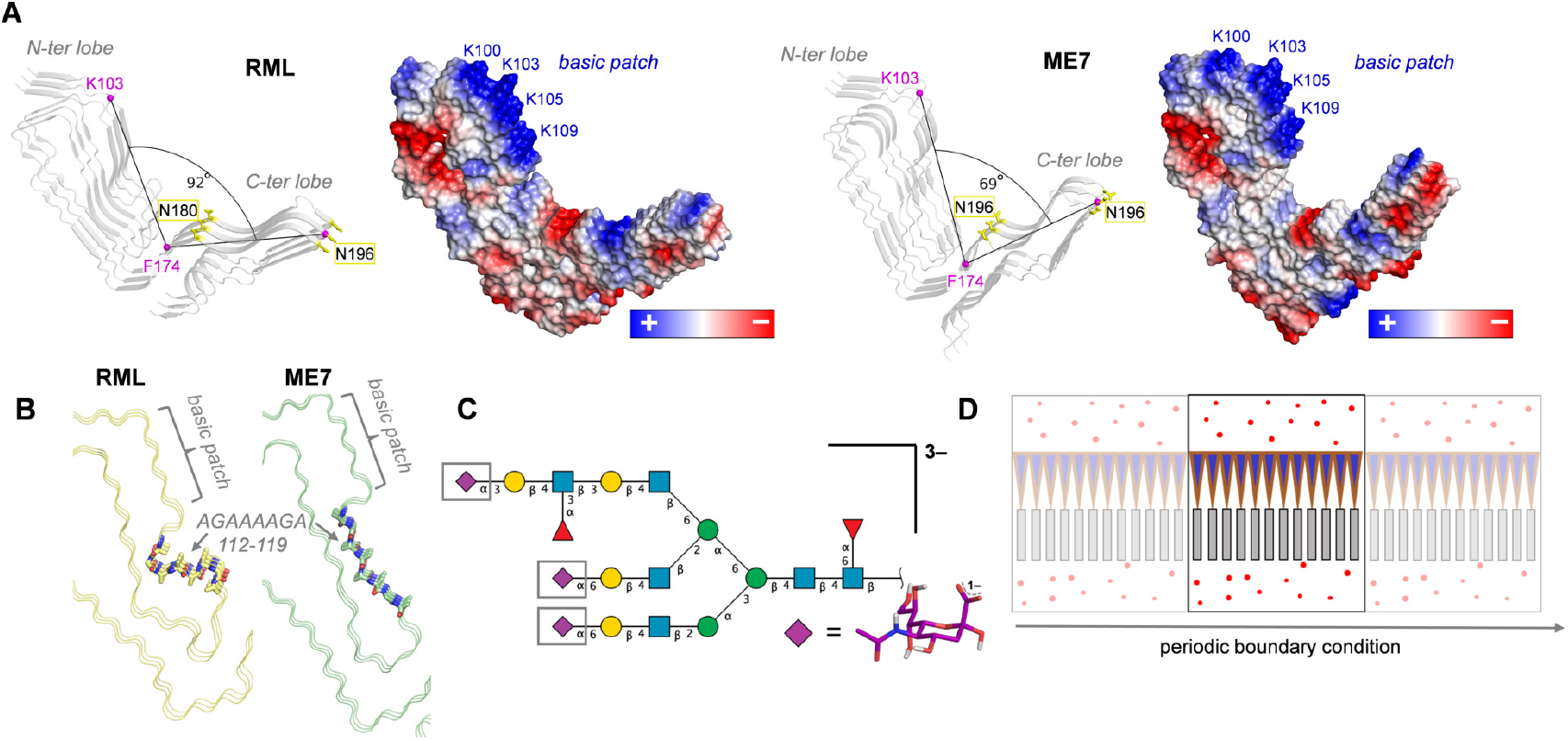
A) Structural representation of the RML and ME7 fibril architectures, deposited under PDB IDs 7QIG^16^ and 8A00,^18^ respectively. For each fibril, a cartoon representation is shown with N180 and N196 highlighted as yellow sticks. The *α*-carbons of K103, F174, and N196 (magenta spheres) are used to define the angle between the N-terminal and C-terminal lobes of the fibril. For each structure, a vacuum electrostatic surface generated with PyMOL’s^22^ protein contact potential is shown, with lysine residues forming the basic patch indicated in blue. B) Conformationally variable region of the RML (yellow) and ME7 (green) fibrils. Amino acids belonging to the AGAAAAGA palindrome (residues 112–119) are shown as sticks, with their spatial relationship to the basic patch indicated. C) Structure of the glycan used for glycosylation at N180 and N196, shown using the symbol nomenclature for glycans: GlcNAc (blue square), Fuc (red triangle), Gal (yellow circle), and Sia (purple diamond). The atomic structure of sialic acid (Sia) is shown as purple sticks. D) Representation of the periodic boundary conditions used to construct the “infinite fibril” model. Fibril strands are depicted as gray rectangles, glycans as purple triangles, and water molecules as red spheres.

Glycans are heterogeneous and highly flexible, and therefore appear only as faint additional densities in cryo-EM maps.^18^ Nevertheless, based on the structural features described above, several hypotheses have been proposed to explain strain-dependent glycoform selectivity:^18^ i) since N180 lies at the bottom of the groove whereas N196 is highly exposed, the glycans at N180 are likely to be more conformationally constrained than those at N196, and thus may play a greater role in determining glycoform selectivity; ii) the short inter-strand spacing in PrP^Sc^ (4.8 Å) enforces a high glycan density, particularly at N180, which may produce steric repulsion opposing fibril propagation; iii) this same dense packing may also induce electrostatic repulsion between negatively charged sialic acid units, further hindering fibril growth; and iv) negatively charged sialic acids may alternatively interact with the basic patch through stabilizing electrostatic contacts.

These considerations alone, however, do not suffice to explain the glycoform selectivities observed in the RML and ME7 strains.^18^ The narrower groove in ME7 would intuitively suggest that N180-linked glycans are more tightly packed than in RML, leading to stronger inter-glycan steric and electrostatic repulsions. This reasoning would predict a lower propensity for ME7 to incorporate diglycosylated forms compared with the wider-grooved RML fibril. In contrast, experimental observations show the opposite trend: ME7 is *more* tolerant to diglycosylated forms than RML.^13^ To date, to the best of our knowledge, only one study has explicitly examined the accommodation of N-linked glycans on a prion fibril model with a parallel in-register *β*-sheet (PIRIBS) architecture using molecular dynamics (MD),^23^ employing an octameric model of truncated human PrP (residues 170–229)^24^ with unsialylated glycans. That work demonstrated that N-linked glycans can be accommodated on successive PIRIBS rungs without destabilizing the octamer over 100 ns of MD simulation.

In the present study, we employ microsecond-scale, all-atom MD simulations under periodic boundary conditions to construct glycosialylated “infinite fibril” models of RML and ME7 fibrils (Figures 1C, 1D, and Methods). This approach eliminates spurious flexibility at the open ends of fibrils and enables detailed characterization of glycan–glycan and glycan–protein interactions. The results allow us to rationalize the experimentally observed glycoform selectivity, addressing an open question as old as the strain phenomenon itself, while establishing a methodology readily applicable to other fibril architectures.

## Methods

### Glycans composition

The high compositional heterogeneity of glycans poses a significant challenge for molecular modeling. Based on available mass spectrometry data,^25,26^ we constructed the representative glycan shown in Figure 1C. This is a triantennary complex-type glycan (bi-, tri-, and tetra-antennary forms have been identified in mouse PrP^26^) featuring core *α*1,6-fucosylation and outer-arm *α*1,3-fucosylation, with a sialyl-Lewis^X^ epitope. It contains three sialic acid residues and thus carries a net charge of –3 at neutral pH, the condition under which the MD simulations were performed. This glycan was used to fully glycosylate the fibril models at both N180 and N196. To assess the specific impact of sialylation, additional MD simulations were carried out using the same glycan with the three sialic acid residues removed.

### Preparation of fibril models for MD

Initial coordinates for the RML and ME7 fibrils were obtained from the electron microscopy structures deposited in the Protein Data Bank under IDs 7QIG and 8A00, respectively. Each structure comprises three strands, and the growth direction of the fibril is aligned with the *z*-axis. Twelve-stranded models were generated by replicating the three-stranded unit three times along the *z*-axis, using an inter-strand spacing of 4.8 Å as determined by cryo-EM.^18^ A three-dimensional model of the sialylated glycan was constructed using the Glycam Carbohydrate Builder^27^ (https://glycam.org/cb/) and downloaded in PDB format. The glycan coordinates were manually aligned to the corresponding N180 and N196 glycosylation sites on the fibril (24 glycans per model). Glycosylated models lacking sialic acid were derived from the sialylated model by removing the sialic acid units. This procedure ensured that the initial glycan geometry was identical in the sialylated and unsialylated systems, thereby minimizing potential bias originating from different starting conformations.

In total, six models were prepared for each strain: unglycosylated, glycosylated with sialylated glycans, and glycosylated with unsialylated glycans. A cubic water box centered on the fibril with a 15 Å buffer from the box borders was generated using AmberTools 24.^28^

### The “infinite fibril” model

A Python script was used to remove water molecules with *z* coordinates exceeding those of the fibril, resulting in twelve-stranded models that are fully solvated along the *x* and *y* directions but unsolvated along *z*, the fibril growth axis. The *z*-dimension of the simulation box was then adjusted to match the fibril length so that periodic boundary conditions accurately reproduce the fibril along the growth direction. This strategy eliminates edge effects and prevents artificial disaggregation caused by finite-size limitations, thereby maintaining fibril stability over long simulation times. The same approach has been previously applied to simulations of amyloid-*β* fibrils.^29,30^

### MD simulations

MD simulations were performed with AMBER 24^28^ using the ff14SB force field^31^ for the PrP protein, the GLYCAM06 force field for the glycans,^32^ and the TIP3P force field for water.^33^ The fibril models were neutralized by adding explicit Na^+^ or Cl^−^ counterions (Li–Merz 12–6 normal usage set^34^).

A two-stage geometry minimization protocol was applied: first, solvent molecules and ions were minimized while keeping the solute fixed; second, a full unrestrained minimization of all atoms in the simulation cell was performed. Equilibration was then carried out through unconstrained MD using a Langevin thermostat^35^ and a Berendsen barostat^36^ with anisotropic pressure scaling. The temperature was increased from 0 to 300 K during the first half of the equilibration and maintained at 300 K thereafter. The SHAKE^37^ algorithm was applied to constrain bonds involving hydrogen, allowing a 2 fs integration time step.

Production simulations were conducted under the same conditions as three independent 500 ns trajectories, yielding a total simulation time of 1.5 *µ*s per system.

### NCIPLOT calculations

Non-covalent interaction (NCI) analysis was carried out using the NCI index as implemented in the NCIPLOT4 program using promolecular charge densities. ^38^ Given a spatial grid around the system of interest, the NCI index identifies regions corresponding to non-covalent interactions through the reduced density gradient *s* (Eq. 1):

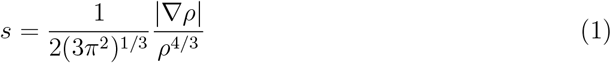

where *ρ* represents the electron density. Non-covalent interactions are detected as regions with simultaneously low *ρ* and low *s* values and are classified according to their position along the *sign*(*λ*_2_)*ρ* scale, where *sign*(*λ*_2_) denotes the sign of the second eigenvalue of the Hessian of *ρ*. Strong attractive interactions appear in the negative region, closed-shell (repulsive) contacts in the positive region, and weak van der Waals interactions near zero. A sample NCIPLOT4 input file is provided in the ESI.

### Analysis

Analysis of the MD trajectories was conducted using `cpptraj`^39^ for hydrogen-bond and total solvent-accessible surface area calculations, and custom Python 3 scripts for strand reordering, root-mean-square fluctuation (RMSF), and per-residue solvent-accessible surface area analyses. The Miller–Madow correction^40^ was applied to estimate the relative entropy of the N180 glycans between the RML and ME7 fibrils, using discrete binning of glycosidic bond torsional angles. Sample analysis scripts are provided in the ESI.

## Results and discussion

### Glycans tune local fibril flexibility

Fully glycosylated 12-mer models of the RML and ME7 fibrils were subjected to 1.5 *µ*s of MD simulation, allowing extensive conformational sampling. Even under full glycosylation (at N180 and N196), glycans could be accommodated without glycan–glycan or glycan–protein steric clashes. We first examined how glycosylation influences local and global fibril flexibility. To assess the global effect of glycosylation on fibril compactness, we computed the distribution of inter-strand distances between equivalent *α*-carbons (C*α*) on consecutive strands for models without glycans, with sialylated glycans, and with glycans lacking sialic acid (Figure 2A).

**Figure 2.**
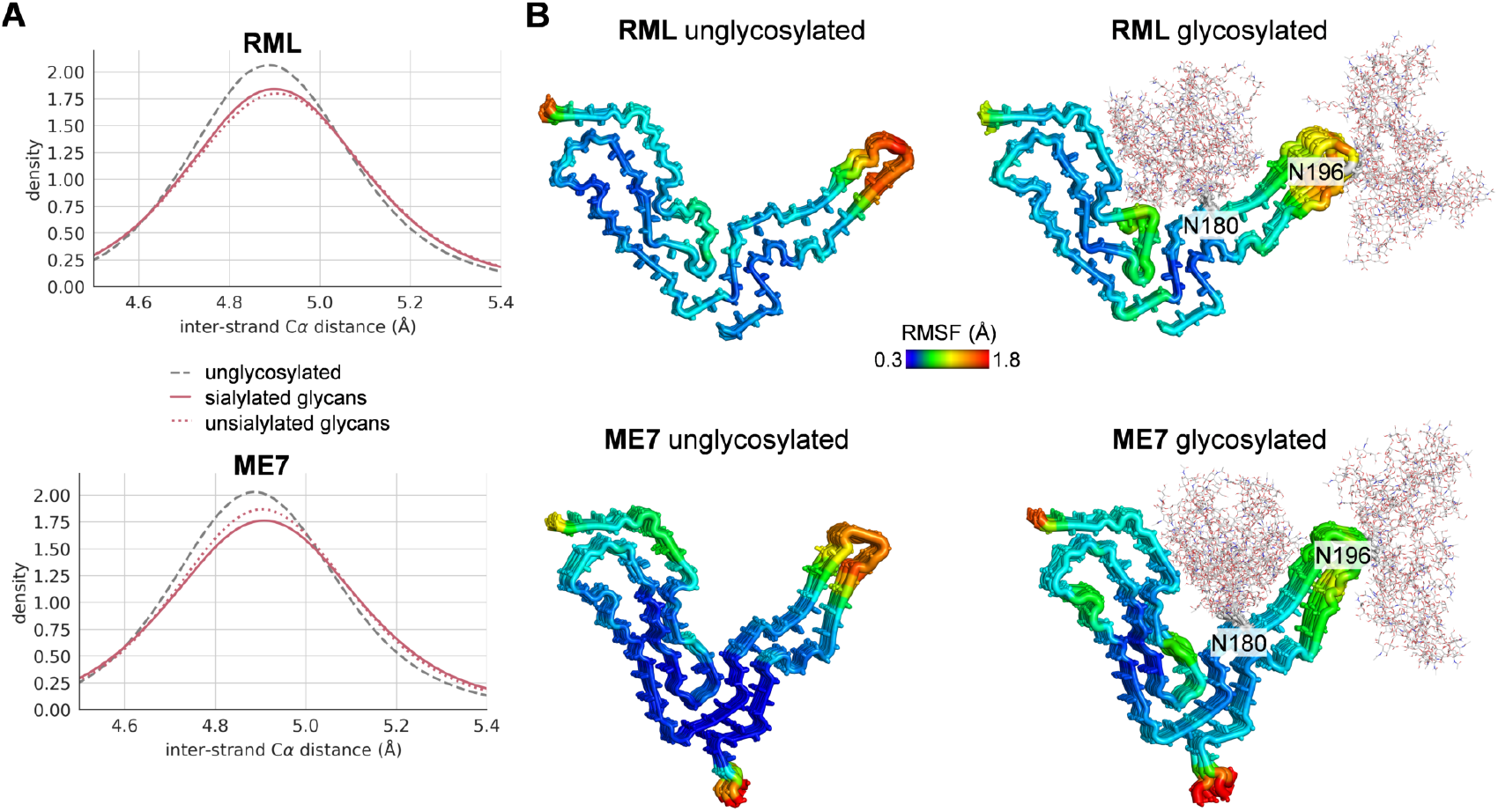
A) Distribution of inter-strand C*α* distances (kernel density estimates) between equivalent residues on consecutive strands from the 1.5 *µ*s MD simulations of RML and ME7 fibrils: unglycosylated (solid grey line), glycosylated with sialylated glycans (solid pink line), and glycosylated with unsialylated glycans (dashed pink line). B) C*α* root-meansquare fluctuations (RMSFs) from the MD simulations of RML and ME7 fibrils, shown for unglycosylated (left) and sialylated glycosylated (right) models. Cartoon thickness and color are scaled to RMSF values, with more rigid structural elements shown as thin blue cartoons and more flexible regions as thick red cartoons. Glycans are depicted as grey lines.

The C*α* distances follow a Gaussian distribution centered at 4.89 Å for the unglycosylated models of both strains, in excellent agreement with the cryo-EM structures, confirming that the fibril models remain stable over long simulation times. Glycosylation with either sialylated or unsialylated glycans induces only a minor effect, slightly broadening the distribution and shifting the maximum by +0.01–0.02 Å toward larger distances. These results demonstrate that glycosylated fibrils are similarly stable, consistent with previous observations on truncated human PrP fibrils.^23^

Second, we assessed the effect of glycosylation on local fibril flexibility by computing the C*α* root-mean-square fluctuations (RMSFs) along the MD trajectories. Figure 2B shows the per-residue C*α* RMSFs for unglycosylated fibrils and for fibrils glycosylated with sialylated glycans, where higher RMSF values indicate greater flexibility. The results show that the fibril core is highly rigid, with the largest flexibility observed at the termini and at the tip of the C-terminal lobe. This finding agrees with the local-resolution map of the RML fibril determined by cryo-EM, thereby validating the model.^16^

N196 glycosylation slightly decreases the flexibility of the C-terminal lobe in both strains, suggesting that lateral packing between glycans on successive strands contributes to structural ordering. Similar trends were observed for the glycosylated models lacking sialic acid (Figure S1).

### Different strain architectures impose distinct conformational constraints on N180 glycans

We next analyzed the flexibility of glycans at the N180 and N196 positions. The per-carbohydrate C1 RMSFs are shown in Figure 3A. While the N196 glycans, which project outward from the C-terminal lobe, are highly flexible in both RML and ME7, the N180 glycans display marked flexibility differences reflecting the distinct groove geometries of the two strains. Specifically, the N180 glycans in ME7 are more conformationally restricted (less flexible) than those in RML. In both cases, the sialic acid residues—highlighted in red in Figure 3A—exhibit the greatest flexibility, consistent with their peripheral position on the outer arm of the glycan and the repulsive interactions between negatively charged units. A similar N180 RMSF difference between RML and ME7 is observed for the unsialylated glycans (Figure S2).

**Figure 3.**
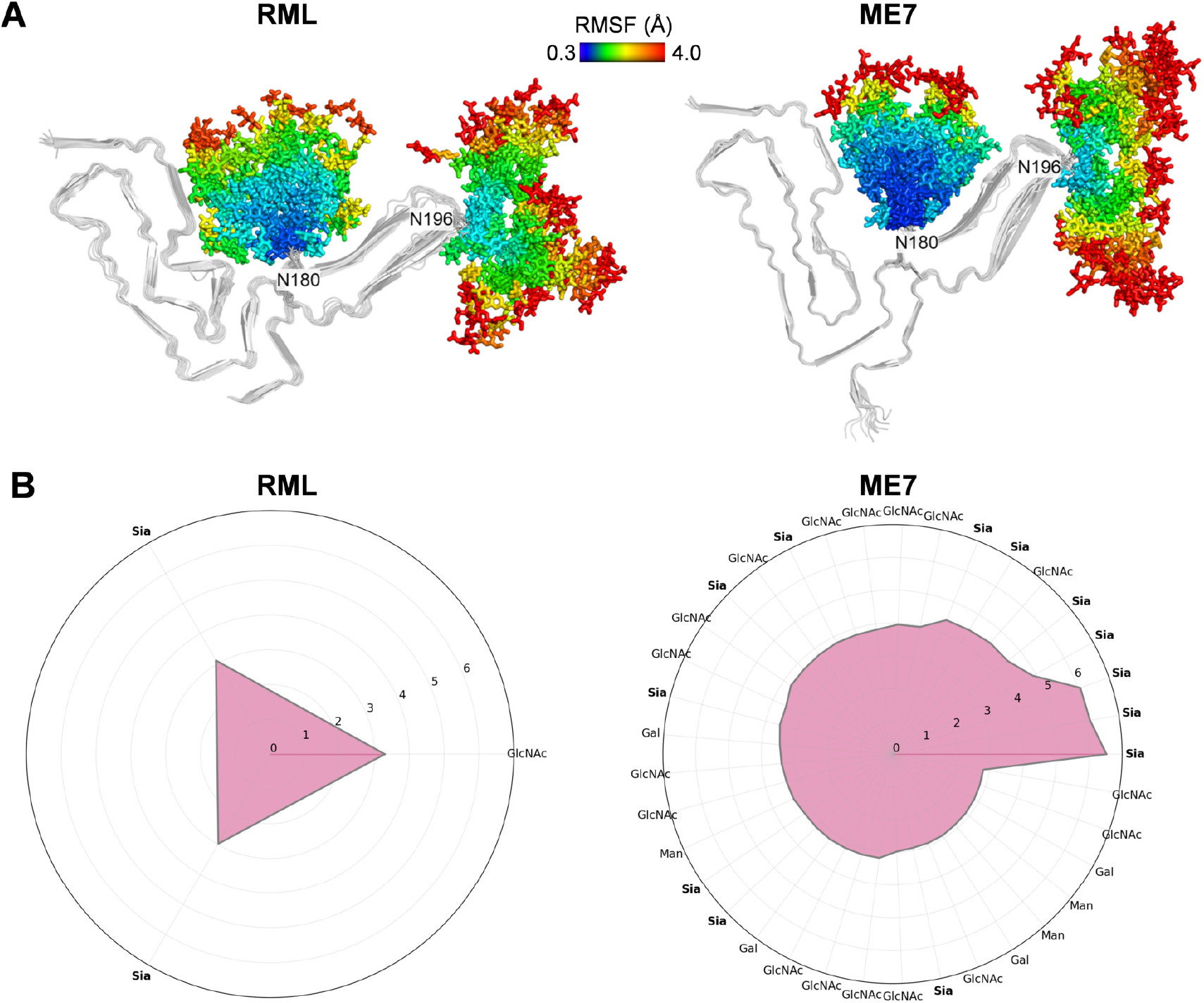
A) Per-carbohydrate RMSFs of the glycans from the 1.5 *µ*s MD simulations of RML and ME7 fibrils glycosylated with sialylated glycans. Fibril strands are shown as grey cartoons, and glycans as sticks. Stick colors are scaled according to RMSF values, with more conformationally constrained carbohydrate units shown in blue and more flexible ones in red. B) Radial plots showing carbohydrate units at position N180 in the RML and ME7 fibrils (glycosylated with sialylated glycans) that participate in cumulative hydrogen bond frequencies of at least 2.8 with other carbohydrate units over the 1.5 *µ*s MD simulations. Because a given carbohydrate unit can form hydrogen bonds with multiple partners simultaneously, cumulative frequency values may exceed 1. Plot tick labels denote carbohydrate types, with sialic acid units (labeled “Sia”) highlighted in bold.

Conformational restriction imposes an entropic penalty, as spatial confinement limits the conformational degrees of freedom of the glycans relative to those in solution. We estimated the relative free-energy difference arising from this conformational confinement between the N180 glycans of RML and ME7 to be approximately 2 kcal mol^−1^ at 300 K (see ESI for further details). This implies that stacking a single glycan at N180 within the ME7 fibril is about 2 kcal mol^−1^ less favorable than stacking the same glycan at the corresponding site in the RML fibril. However, tighter glycan packing also increases the likelihood of forming stabilizing glycan–glycan hydrogen bonds (typically in the 1–7 kcal mol^−1^ range), which can yield favorable enthalpic contributions that partially or fully compensate for the unfavorable entropic term.

Hydrogen bond frequencies between sialylated N180 glycans are shown in Figure 3B, which lists the carbohydrate units involved in at least 2.8 hydrogen bonds with other glycans throughout the MD simulation (corresponding plots for unsialylated glycans are shown in Figure S3). Glycans form a much denser hydrogen-bond network in ME7 than in RML (38 vs 3), as glycan compaction within the narrower groove promotes glycan–glycan interactions. In ME7, the sialic acid units (highlighted in bold in Figure 3B) establish highly persistent hydrogen bonds that partially shield the negative charge, thereby reducing electrostatic repulsion. This denser hydrogen-bond network in ME7 at least partially compensates for the higher entropic penalty associated with conformational restrictions, consistent with the well-known entropy–enthalpy compensation phenomenon.^41^ These findings suggest that glycan–glycan interactions alone do not determine the glycoform preference of the strains; rather, glycans can be accommodated without steric clashes and can engage in favorable glycan–glycan interactions. Thus, the origin of glycoform preferences likely lies in glycan–fibril interactions.

### Glycans engage prion fibrils in a wide range of non-covalent interactions

Glycan–fibril contacts involve several types of non-covalent interactions (NCIs), which engage distinct regions of the protein surface depending on the strain. Figure 4 shows an overlay of thirty frames extracted from the MD simulations of the two glycosialylated fibrils. As discussed above, the glycans exhibit high mobility and therefore sample a broad conformational space. While the N196 glycan remains largely solvent-exposed throughout the trajectory, the N180 glycan forms numerous contacts involving both the N-terminal and C-terminal sides of the groove.

**Figure 4.**
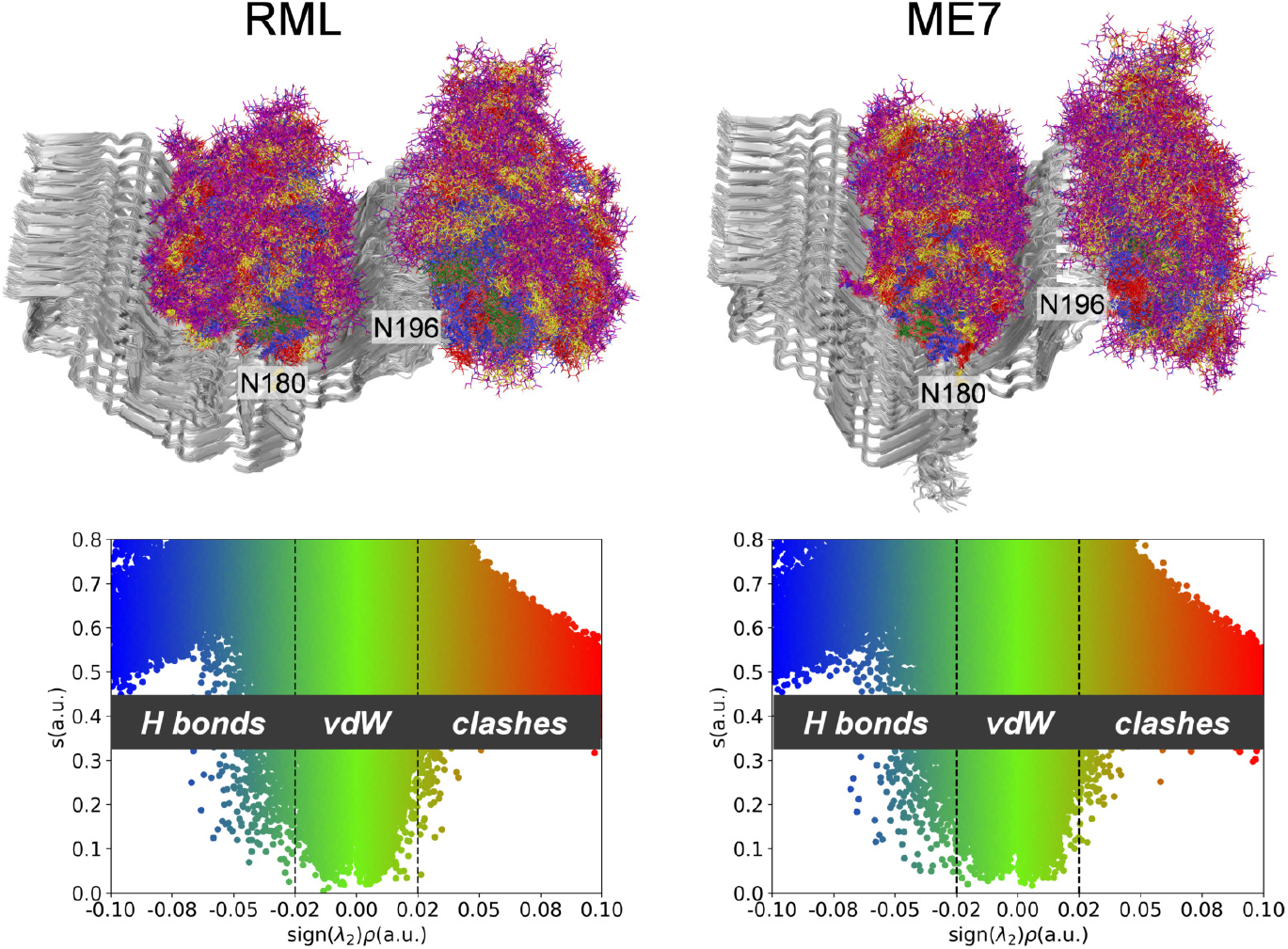
Top: Overlay of thirty evenly spaced frames extracted from the 1.5 *µ*s MD simulations of RML and ME7 fibrils glycosylated with sialylated glycans. Fibril strands are depicted as grey cartoons, and glycans as lines. GlcNAc units are shown in blue, Fuc in red, Gal in yellow, and Sia in purple. Bottom: Two-dimensional plots of the low *ρ* (electron density) and low *s* (reduced density gradient) regions corresponding to non-covalent interactions, computed from the same 30 frames. Strong attractive interactions appear in the negative region, closed-shell (repulsive) contacts in the positive region, and weak van der Waals interactions near zero. Vertical lines at an arbitrary value of 0.02 are included as visual guides to distinguish between different interaction types.

To characterize the nature of these glycan–fibril contacts, we performed a non-covalent interaction analysis along the MD trajectories using the NCI index (see Methods). This approach provides a qualitative, interaction-type–agnostic assessment of all NCIs in the system and has been applied in the study of non-covalent contacts in biomolecular systems, including protein fibrils.^42,43^

The resulting plots are shown in Figure 4. Points in the lower region of the plots (low *s* values, where *s* is the reduced density gradient) correspond to non-covalent interactions. Their position along the *x*-axis (*sign*(*λ*_2_)*ρ*) reflects both the nature and strength of the interactions: strong attractive contacts, such as hydrogen bonds, occur at negative values (blue); closed-shell interactions, including steric repulsion, appear at positive values (red); and weak van der Waals contacts cluster near zero (green). Both strains exhibit overall similar NCI distributions, dominated by weak van der Waals interactions, with a certain proportion of attractive interactions (hydrogen bonds) and no significant steric clashes.

On one hand, the affinity of the basic patch for negatively charged species suggests that hydrogen bonds between lysine residues and sialic acid units may contribute to the overall stabilization of the system. On the other hand, glycans can shield the fibril surface, potentially occluding the open ends of the growing fibril and thereby interfering with the aggregation process. To elucidate the origin of the distinct glycoform selectivities observed in RML and ME7, we quantified both effects.

### Sialylation at N180 strengthens hydrogen bonding to the basic patch in ME7 relative to RML

Hydrogen bond frequencies between glycans and fibrils are summarized in Figure 5, which lists the amino acid residues engaged in at least one hydrogen bond for at least half of the MD trajectory. For each residue, the cumulative hydrogen bond fraction is decomposed into contributions from the different carbohydrate types. In both strains, lysine residues (in bold) within the basic patch form persistent hydrogen bonds with the glycans, with sialic acid accounting for approximately half of these interactions.

**Figure 5.**
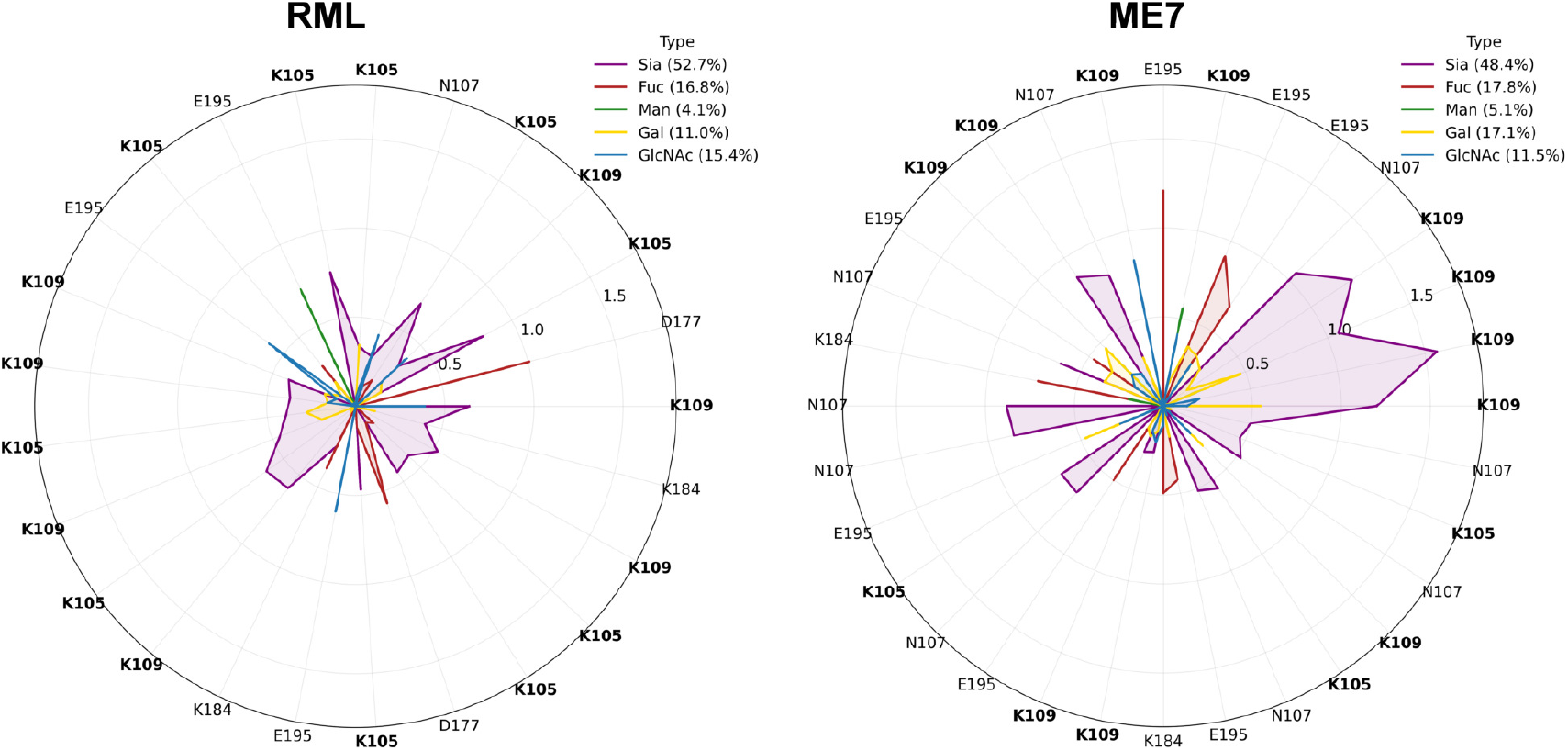
Radial plots showing amino acid residues from fibril models glycosylated with sialylated glycans that participate in cumulative hydrogen bond frequencies with glycans of at least 0.5 over the 1.5 *µ*s MD simulations. Cumulative hydrogen bond frequencies are decomposed into contributions from individual carbohydrate types: GlcNAc (blue), Fuc (red), Gal (yellow), and Sia (purple). Because a single amino acid can simultaneously form hydrogen bonds with multiple carbohydrate units, cumulative frequency values may exceed 1. Plot tick labels indicate the amino acid identities, with lysines (K) highlighted in bold. The percentage contribution of each carbohydrate type to the total glycan–fibril hydrogen bonds is shown.

These hydrogen bonds are slightly more frequent in ME7 (32) than in RML (25) and primarily involve lysines K105 and K109. Notably, the ME7 strain exhibits more persistent (stronger) interactions between sialic acid and K109 compared to the RML strain (Figures 5 and 6A).

**Figure 6.**
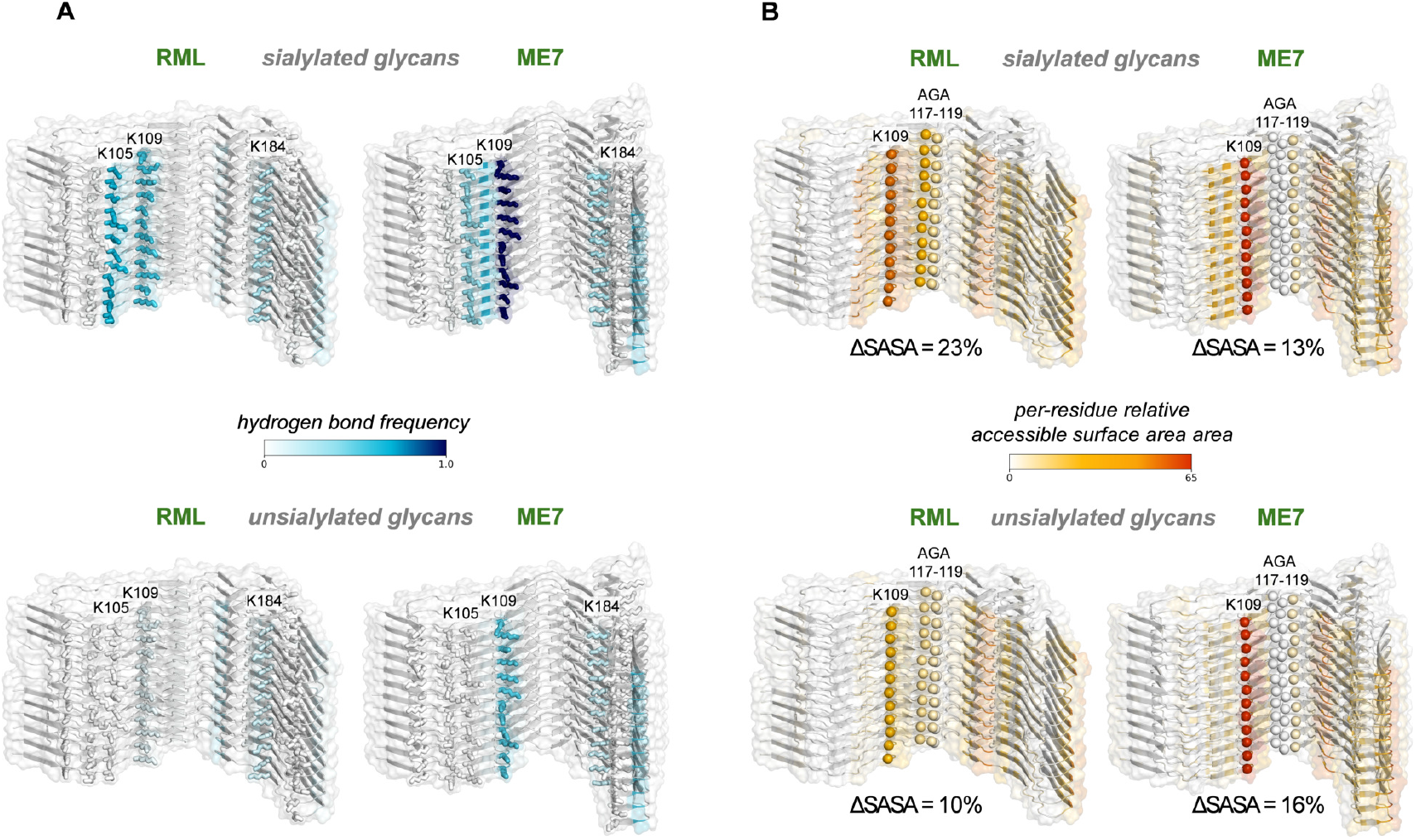
A) Per-residue cumulative frequency of glycan–fibril hydrogen bonds mapped onto the three-dimensional structures of RML and ME7 fibrils glycosylated with sialylated glycans (top) and unsialylated glycans (bottom). The fibril backbone is shown as a cartoon, and lysine side chains are represented as sticks. Structure coloring corresponds to hydrogen bond frequency, with residues showing the lowest frequencies in white and those with the highest frequencies in blue. Frequencies are averaged per residue over the 12 strands composing each model. B) Per-residue differences in solvent-accessible surface area (SASA) between unglycosylated RML and ME7 fibril models and their corresponding glycosylated counterparts with sialylated glycans (top) and unsialylated glycans (bottom). The fibril backbone is shown as a cartoon. The C*α* atoms of K109 and of the AGA 117–119 segment are shown as spheres. Structure coloring corresponds to the difference in per-residue SASA, representing the degree of shielding conferred by glycans: residues with minimal shielding are shown in white, and those with maximal shielding in red. Per-residue SASA differences are averaged over the 12 strands of each model. The global shielding provided by glycans to the whole fibril is indicated below each model as the percentage of total surface area (ΔSASA).

Because interactions between oppositely charged groups are expected to enhance overall system stability, this difference—arising from the relative spatial arrangement of the glycans and the basic patch, which is dictated by the fibril architecture—constitutes a key structural distinction between the two strains. It provides a plausible partial explanation for the higher tolerance of ME7 toward diglycosylated forms: since the same glycan forms more effective stabilizing interactions in ME7 than in RML, diglycosylated glycoforms are likely subject to weaker negative selection for incorporation into the fibril in ME7.

For unsialylated glycans, hydrogen bond frequencies are generally lower, and all persistent hydrogen bonds involving the basic patch are lost in the RML fibril, while in the ME7 fibril they decrease from 14 to 6 (Figure S4). This confirms that electrostatic interactions between opposite charges are a major driving force for glycan–basic patch contacts.

### Sialylated glycans provide extensive shielding and specific engagement of the palindrome sequence in RML but not in ME7

To assess the overall extent of glycan–fibril contacts, we calculated the solvent-accessible surface area (SASA, see Methods) along the MD trajectories for both glycosylated and unglycosylated fibrils, and used these values to determine the fraction of the fibril surface shielded by glycans. For sialylated glycans, we observed a pronounced difference between the two strains: 23% shielding in RML versus only 13% in ME7. This indicates that, in the RML fibril, the same glycans not only provide weaker electrostatic compensation for the basic patch but also impose a greater overall shielding effect than in ME7.

To evaluate the local distribution of this shielding, we computed the per-residue SASA along the MD simulations and mapped onto the structures the differences between the unglycosylated and glycosylated models (Figure 6B). For each residue, a large SASA difference indicates strong shielding by glycans (i.e., reduced solvent accessibility upon glycosylation), whereas small differences correspond to minimal or no shielding. The groove between the N-terminal and C-terminal lobes exhibits the strongest shielding, consistent with the spatial confinement of the N180 glycan, which maintains extensive contacts with the fibril. Unexpectedly, however, the broader groove of the RML fibril is more shielded than the narrower one of ME7.

This arises from differences in the conformation of the variable region that contains the palindrome sequence (Figure 1B). In RML, the second half of this stretch protrudes toward the groove center, whereas in ME7 it is recessed behind the basic patch. Consequently, this conformational disparity produces distinct shielding patterns of this region—which is critical for prion aggregation—between the two strains: in RML, the AGA 117–119 residues are substantially shielded by glycans (Figure 6B), whereas in ME7 they remain largely exposed.

Taken together, these findings reveal a second structural basis for the lower propensity of RML to accommodate diglycosylated glycoforms: the same N180 glycans exert stronger shielding in RML than in ME7 and specifically interact with the palindrome sequence, which remains unengaged in ME7. Thus, in RML, the glycan not only occludes a larger fibril surface but may also directly interfere with a region essential for propagation.

Removal of sialic acids markedly alters both global and per-residue SASA differences in RML, reducing overall shielding from 23% to 10% and diminishing engagement of the AGA 117–119 residues (Figure 6B). This correlates with the loss of hydrogen bonds involving the basic patch (Figure 6A) and demonstrates that, in the absence of the electrostatic attraction between lysines and sialic acids, glycans form fewer contacts with the fibril in this strain. In contrast, desialylation has a smaller effect in ME7: sialylated and unsialylated glycans confer comparable global shielding (13% vs. 16%, respectively; Figure 6B), the AGA 117–119 residues remain accessible, and K109 continues to participate in stable hydrogen bonds (Figure 6A).

Overall, the analysis of glycan–fibril contacts supports a threefold hypothesis explaining the sialoglycoform preferences of the RML and ME7 strains, based on distinct interactions involving the N180 glycan. With identical glycosylation patterns, the RML fibril (i) exhibits greater overall surface shielding than ME7, and (ii) only in RML do the glycans directly contact the palindrome sequence. Since such interactions are expected to hinder the recruitment of additional monomeric units, RML displays lower tolerance to diglycosylated forms, likely to avoid excessive accumulation of consecutive glycans at N180. Considering not only the extent but also the nature of the contacts, (iii) RML is further disadvantaged relative to ME7 by weaker stabilizing interactions between sialoglycans and the basic patch—again consistent with reduced tolerance to glycosylation at N180.

Regarding specifically the role of sialylation, our simulations show that RML is more sensitive than ME7 to the presence of sialic acid, as its inclusion or removal more strongly alters glycan–fibril interactions. This observation aligns with the experimentally reported differential behavior of the two strains with respect to sialylation.^13^ In both strains, the abundances of diglycosylated and hypersialylated glycoforms are anticorrelated; however, desialylation increases the proportion of diglycosylated glycoforms by only ∼10% in ME7, but by ∼20% in RML.^13^

Although modeling complex glycoprotein aggregates remains inherently limited by the heterogeneity of glycosylation patterns, the present simulations combine extensive conformational sampling with a rigorous approach for modeling the periodic structure of prion fibrils. This enables the derivation of detailed structural insights into the molecular origins of the glycoform selectivities observed in the RML and ME7 strains.

## Conclusions

We combined high-resolution cryo-EM structures with microsecond-scale molecular dynamics simulations under periodic boundary conditions to characterize in detail the effects of different glycosylation patterns on the mouse PrP strains RML and ME7. Our results show that sialylation is a key determinant of the extent of glycan–fibril contacts through specific hydrogen-bond interactions with the basic patch. These interactions are stronger in ME7 than in RML, despite the lower overall fibril surface shielding observed in ME7. We also show that only in the RML fibril do the N180 glycans have the potential to occlude the AGAAAAGA palindrome sequence, which is essential for prion propagation. Together, these findings account for (i) the lower tolerance of RML to diglycosylated forms compared with ME7 and (ii) the greater change in glycoform ratios upon desialylation observed in RML relative to ME7.

This work establishes a general methodology for characterizing glycan–fibril interactions, providing a framework to obtain structural insights into the origins of glycoform preferences in other strains with known fibril architectures and to assess how glycan shielding may interfere with the binding of candidate aggregation inhibitors. The same approach can also be applied to explore additional aspects of prion aggregation, such as the preferred arrangement of glycoforms for a given di-, mono- and unglycosylated ratio (e.g., stoichiometric versus probabilistic assembly), and to elucidate the individual effect of the two glycosylation positions as well as that of the GPI anchor on fibril properties.

## Acknowledgement

This research has been funded by MCIN/AEI/10.13039/501100011033 (grants RYC2022-036457-I and EUR2023-143462). FP thanks Prof. Gonzalo Jiménez-Osés, Prof. Joaquín Castilla and Dr. Hasier Eraña for fruitful discussions.

## Supporting Information Available

C*α* and per-carbohydrate-unit root-mean-square fluctuations from the MD simulations of RML and ME7 fibrils glycosylated with unsialylated glycans. Radial plots showing carbohydrate units at position N180 in the RML and ME7 fibrils glycosylated with unsialylated glycans that participate in glycan-glycan hydrogen bonds. Radial plots showing amino acids of the RML and ME7 fibrils glycosylated with unsialylated glycans that participate in glycan-fibril hydrogen bonds. Python 3 script used for estimating glycans torsional entropies. Sample input employed for NCIPLOT4 calculations. Sample Python 3 scripts used for analysis: strand reordering, fibril and glycan root-mean-square fluctuations calculations, per-residue solvent accessible surface area. The following data are available through the Zenodo repository https://zenodo.org/records/17512164: MD trajectories for unglycosylated fibrils, fibrils glycosylated with sialylated glycans and fibrils glycosylated with unsialylated glycans (PyMOL sessions, PyMOL open source code can be downloaded from https://github.com/schrodinger/pymol-open-source).

## TOC Graphic

## References

(1) Tuzi, N. L.; Cancellotti, E.; Baybutt, H.; Blackford, L.; Bradford, B.; Plinston, C.; Coghill, A.; Hart, P.; Piccardo, P.; Barron, R. M.; Manson, J. C. Host PrP Glycosylation: A Major Factor Determining the Outcome of Prion Infection. PLOS Biol. 2008, 6, 1–11.

(2) Wiseman, F. K.; Cancellotti, E.; Piccardo, P.; Iremonger, K.; Boyle, A.; Brown, D.; Ironside, J. W.; Manson, J. C.; Diack, A. B. The Glycosylation Status of PrP^C^ Is a Key Factor in Determining Transmissible Spongiform Encephalopathy Transmission between Species. J. Virol. 2015, 89, 4738–4747.

(3) Johnson, R. T. Prion diseases. Lancet Neurol. 2005, 4, 635–642.

(4) Cobb, N. J.; Surewicz, W. K. Prion Diseases and Their Biochemical Mechanisms. Biochemistry 2009, 48, 2574–2585.

(5) Walker, L. C.; Jucker, M. The prion principle and Alzheimer’s disease. Science 2024, 385, 1278–1279.

(6) Tanaka, M.; Collins, S. R.; Toyama, B. H.; Weissman, J. S. The physical basis of how prion conformations determine strain phenotypes. Nature 2006, 442, 585–589.

(7) Morales, R.; Abid, K.; Soto, C. The prion strain phenomenon: Molecular basis and unprecedented features. Biochim. Biophys. Acta - Mol. Basis Dis. 2007, 1772, 681– 691.

(8) Aguzzi, A.; Heikenwalder, M.; Polymenidou, M. Insights into prion strains and neurotoxicity. Nat. Rev. Mol. Cell Biol. 2007, 8, 552–561.

(9) Aguzzi, A. Unraveling prion strains with cell biology and organic chemistry. Proc. Natl. Acad. Sci. U.S.A 2008, 105, 11–12.

(10) Igel, A.; Fornara, B.; Rezaei, H.; Béringue, V. Prion assemblies: structural heterogeneity, mechanisms of formation, and role in species barrier. Cell Tissue Res. 2023, 392, 149–166.

(11) Khalili-Shirazi, A.; Summers, L.; Linehan, J.; Mallinson, G.; Anstee, D.; Hawke, S.; Jackson, G. S.; Collinge, J. PrP glycoforms are associated in a strain-specific ratio in native PrPSc. J. Gen. Virol. 2005, 86, 2635–2644.

(12) Xanthopoulos, K.; Polymenidou, M.; Bellworthy, S. J.; Benestad, S. L.; Sklaviadis, T. Species and Strain Glycosylation Patterns of PrPSc. PLOS ONE 2009, 4, 1–10.

(13) Katorcha, E.; Makarava, N.; Savtchenko, R.; Baskakov, I. V. Sialylation of the prion protein glycans controls prion replication rate and glycoform ratio. Sci. Rep. 2015, 5, 16912.

(14) Katorcha, E.; Makarava, N.; Savtchenko, R.; d’Azzo, A.; Baskakov, I. V. Sialylation of Prion Protein Controls the Rate of Prion Amplification, the Cross-Species Barrier, the Ratio of PrPSc Glycoform and Prion Infectivity. PLOS Pathog. 2014, 10, 1–12.

(15) Kraus, A.; Hoyt, F.; Schwartz, C. L.; Hansen, B.; Artikis, E.; Hughson, A. G.; Raymond, G. J.; Race, B.; Baron, G. S.; Caughey, B. High-resolution structure and strain comparison of infectious mammalian prions. Mol. Cell 2021, 81, 4540–4551.e6.

(16) Manka, S. W.; Zhang, W.; Wenborn, A.; Betts, J.; Joiner, S.; Saibil, H. R.; Collinge, J.; Wadsworth, J. D. F. 2.7 Å cryo-EM structure of ex vivo RML prion fibrils. Nat. Commun. 2022, 13, 4004.

(17) Hoyt, F.; Standke, H. G.; Artikis, E.; Schwartz, C. L.; Hansen, B.; Li, K.; Hughson, A. G.; Manca, M.; Thomas, O. R.; Raymond, G. J.; Race, B.; Baron, G. S.; Caughey, B.; Kraus, A. Cryo-EM structure of anchorless RML prion reveals variations in shared motifs between distinct strains. Nat. Commun. 2022, 13, 4005.

(18) Manka, S. W.; Wenborn, A.; Betts, J.; Joiner, S.; Saibil, H. R.; Collinge, J.; Wadsworth, J. D. F. A structural basis for prion strain diversity. Nat. Chem. Biol. 2023, 19, 607–613.

(19) Hoyt, F.; Alam, P.; Artikis, E.; Schwartz, C. L.; Hughson, A. G.; Race, B.; Baune, C.; Raymond, G. J.; Baron, G. S.; Kraus, A.; Caughey, B. Cryo-EM of prion strains from the same genotype of host identifies conformational determinants. PLOS Pathog. 2022, 18, 1–19.

(20) Alam, P.; Hoyt, F.; Artikis, E.; Soukup, J.; Hughson, A. G.; Schwartz, C. L.; Barbian, K.; Miller, M. W.; Race, B.; Caughey, B. Cryo-EM structure of a natural prion: chronic wasting disease fibrils from deer. Acta Neuropathol. 2024, 148, 56.

(21) Norstrom, E. M.; Mastrianni, J. A. The AGAAAAGA Palindrome in PrP Is Required to Generate a Productive PrP^Sc^-PrP^C^ Complex That Leads to Prion Propagation. J. Biol. Chem. 2005, 280, 27236–27243.

(22) Schrödinger, LLC, The PyMOL Molecular Graphics System, Version 1.8. 2015,

(23) Artikis, E.; Roy, A.; Verli, H.; Cordeiro, Y.; Caughey, B. Accommodation of In-Register N-Linked Glycans on Prion Protein Amyloid Cores. ACS Chem. Neurosci. 2020, 11, 4092–4097.

(24) Wang, L.-Q.; Zhao, K.; Yuan, H.-Y.; Wang, Q.; Guan, Z.; Tao, J.; Li, X.-N.; Sun, Y.; Yi, C.-W.; Chen, J.; Li, D.; Zhang, D.; Yin, P.; Liu, C.; Liang, Y. Cryo-EM structure of an amyloid fibril formed by full-length human prion protein. Nat. Struct. Mol. Biol. 2020, 27, 598–602.

(25) Rudd, P. M.; Endo, T.; Colominas, C.; Groth, D.; Wheeler, S. F.; Harvey, D. J.; Wormald, M. R.; Serban, H.; Prusiner, S. B.; Kobata, A.; Dwek, R. A. Glycosylation differences between the normal and pathogenic prion protein isoforms. Proc. Natl. Acad. Sci. USA 1999, 96, 13044–13049.

(26) Stimson, E.; Hope, J.; Chong, A.; Burlingame, A. L. Site-Specific Characterization of the N-Linked Glycans of Murine Prion Protein by High-Performance Liquid Chromatography/Electrospray Mass Spectrometry and Exoglycosidase Digestions. Biochemistry 1999, 38, 4885–4895.

(27) Grant, O. C.; Wentworth, D.; Holmes, S. G.; Kandel, R.; Sehnal, D.; Wang, X.; Xiao, Y.; Sheppard, P.; Grelsson, T.; Coulter, A.; Miller, G.; Foley, B. L.; Woods, R. J. Generating 3D Models of Carbohydrates with GLYCAM-Web. bioRxiv, doi: 10.1101/2025.05.08.652828 2025,

(28) Case, D. et al. Amber 2025. 2025.

(29) Buchete, N.-V.; Tycko, R.; Hummer, G. Molecular Dynamics Simulations of Alzheimer’s β-Amyloid Protofilaments. J. Mol. Biol. 2005, 353, 804–821.

(30) Tywoniuk, B.; Yuan, Y.; McCartan, S.; Szydlowska, B. M.; Tofoleanu, F.; Brooks, B. R.; Buchete, N.-V. Amyloid Fibril Design: Limiting Structural Polymorphism in Alzheimer’s Aβ Protofilaments. J. Phys. Chem. B 2018, 122, 11535–11545.

(31) Maier, J. A.; Martinez, C.; Kasavajhala, K.; Wickstrom, L.; Hauser, K. E.; Simmerling, C. ff14SB: Improving the Accuracy of Protein Side Chain and Backbone Parameters from ff99SB. J. Chem. Theory Comput. 2015, 11, 3696–3713.

(32) Kirschner, K. N.; Yongye, A. B.; Tschampel, S. M.; González-Outeiriño, J.; Daniels, C. R.; Foley, B. L.; Woods, R. J. GLYCAM06: A generalizable biomolecular force field. Carbohydrates. J. Comput. Chem. 2008, 29, 622–655.

(33) Jorgensen, W. L.; Chandrasekhar, J.; Madura, J. D.; Impey, R. W.; Klein, M. L. Comparison of simple potential functions for simulating liquid water. J. Chem. Phys. 1983, 79, 926–935.

(34) Sengupta, A.; Li, Z.; Song, L. F.; Li, P.; Merz, K. M. J. Parameterization of Monovalent Ions for the OPC3, OPC, TIP3P-FB, and TIP4P-FB Water Models. J. Chem. Inf. Model. 2021, 61, 869–880.

(35) Loncharich, R. J.; Brooks, B. R.; Pastor, R. W. Langevin dynamics of peptides: The frictional dependence of isomerization rates of N-acetylalanyl-N’-methylamide. Biopolymers 1992, 32, 523–535.

(36) Berendsen, H. J. C.; Postma, J. P. M.; van Gunsteren, W. F.; DiNola, A.; Haak, J. R. Molecular dynamics with coupling to an external bath. J. Chem. Phys. 1984, 81, 3684– 3690.

(37) Miyamoto, S.; Kollman, P. A. Settle: An analytical version of the SHAKE and RATTLE algorithm for rigid water models. J. Comput. Chem. 1992, 13, 952–962.

(38) Boto, R. A.; Peccati, F.; Laplaza, R.; Quan, C.; Carbone, A.; Piquemal, J.-P.; Maday, Y.; Contreras-Garcia, J. NCIPLOT4: Fast, Robust, and Quantitative Analysis of Noncovalent Interactions. J. Chem. Theory Comput. 2020, 16, 4150–4158.

(39) Roe, D. R.; Cheatham, T. E. I. PTRAJ and CPPTRAJ: Software for Processing and Analysis of Molecular Dynamics Trajectory Data. J. Chem. Theory Comput. 2013, 9, 3084–3095.

(40) Miller, G. Note on the bias of information estimates. Info. Theory Psychol. Prob. Methods 1955, II-B, 4005.

(41) Peccati, F.; Jiménez-Osés, G. Enthalpy–Entropy Compensation in Biomolecular Recognition: A Computational Perspective. ACS Omega 2021, 6, 11122–11130.

(42) Peccati, F. NCIPLOT4 Guide for Biomolecules: An Analysis Tool for Noncovalent Interactions. J. Chem. Inf. Model 2020, 60, 6–10.

(43) Urriolabeitia, A.; Contreras-García, J.; De Sancho, D.; López, X. Resolving Molecular Interactions in Protein Folding Trajectories with NCIPLOT. J. Chem. Inf. Model 2025, 65, 10613–10623.

