## Supplementary information for "Structural Basis of Glycoform Selectivity in Prion Strains"

Francesca Peccati*,†,‡

**3 Sample NCIPLOT4 input** **25**

**4 Sample script for strand reordering** **25**

**5 Sample script for C$\alpha$ RMSF calculation** **29**

**6 Sample script for glycans RMSF calculation** **30**

**7 Sample script for per-residue SASA analysis** **32**

### 1 Supplementary figures

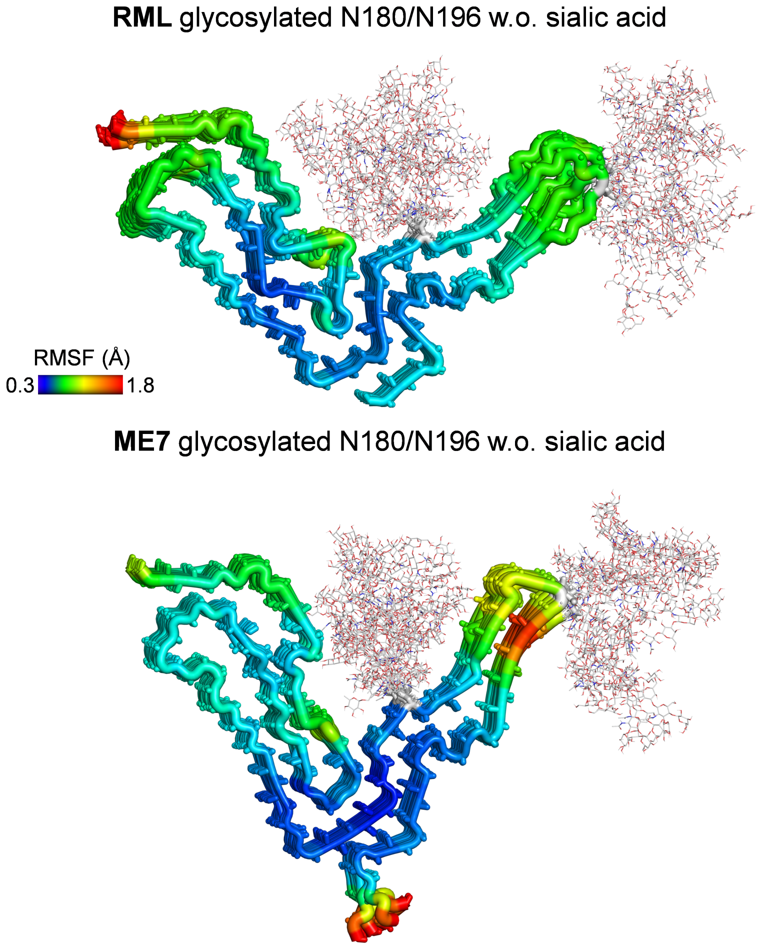

Figure S 1: C$\alpha$ root-mean-square fluctuations (RMSFs) from the MD simulations of RML and ME7 fibrils glycosylated with unsialylated glycans. Cartoon thickness and color are scaled to RMSF values, with more rigid structural elements shown as thin blue cartoons and more flexible regions as thick red cartoons. Glycans are depicted as grey lines.

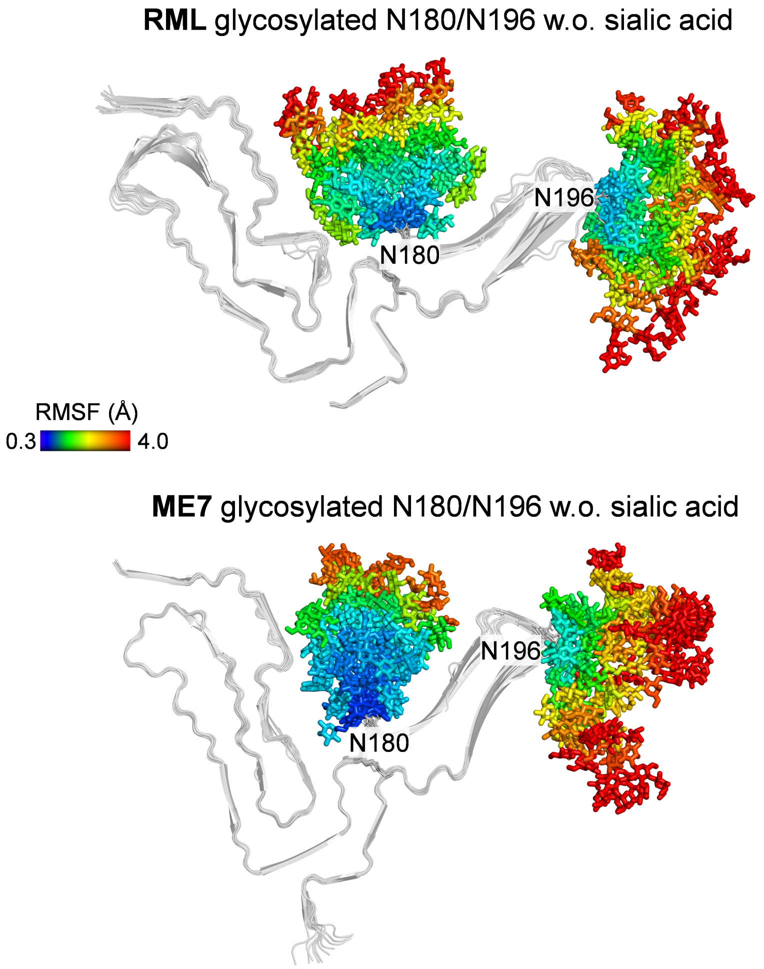

Figure S 2: Per-carbohydrate RMSFs of the glycans from the 1.5 $\mu$s MD simulations of RML and ME7 fibrils glycosylated with unsialylated glycans. Fibril strands are shown as grey cartoons, and glycans as sticks. Stick colors are scaled according to RMSF values, with more conformationally constrained carbohydrate units shown in blue and more flexible ones in red.

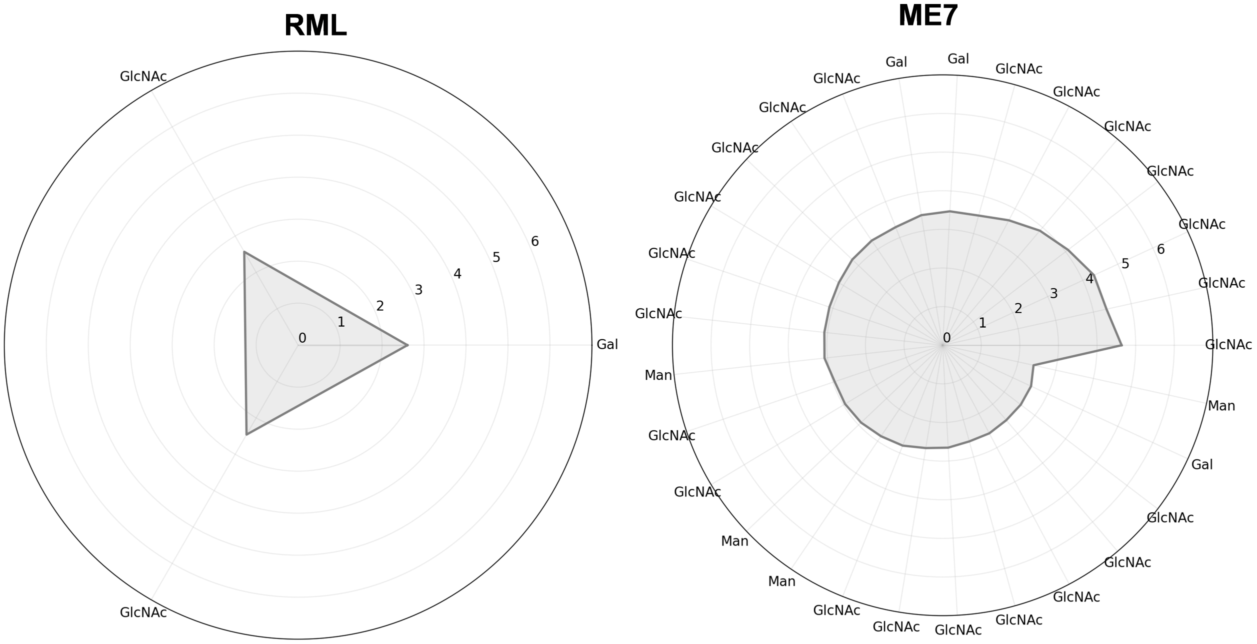

Figure S 3: Radial plots showing carbohydrate units at position N180 in the RML and ME7 fibrils (glycosylated with unsialylated glycans) that participate in cumulative hydrogen bond frequencies of at least 2.4 with other carbohydrate units over the 1.5 $\mu$s MD simulations. Because a given carbohydrate unit can form hydrogen bonds with multiple partners simultaneously, cumulative frequency values may exceed 1. Plot tick labels denote carbohydrate types.

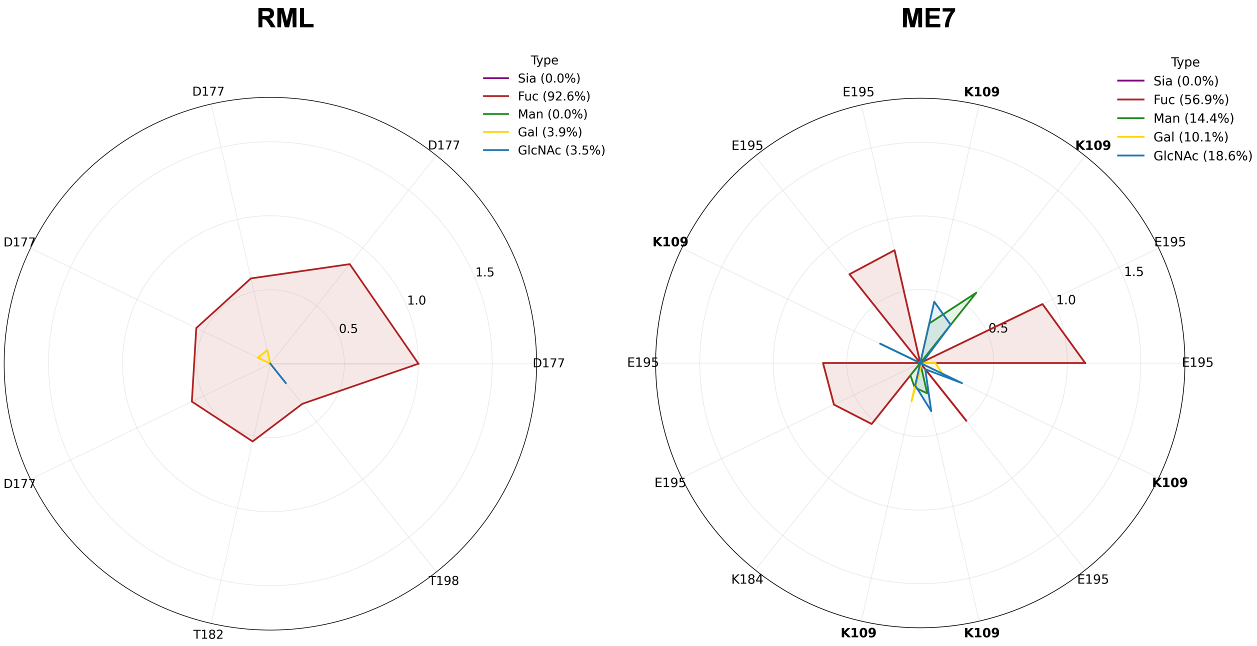

Figure S 4: Radial plots showing amino acid residues from fibril models glycosylated with unsialylated glycans that participate in cumulative hydrogen bond frequencies with glycans of at least 0.5 over the 1.5 $\mu$s MD simulations. Cumulative hydrogen bond frequencies are decomposed into contributions from individual carbohydrate types: GlcNAc (blue), Fuc (red) and Gal (yellow). Because a single amino acid can simultaneously form hydrogen bonds with multiple carbohydrate units, cumulative frequency values may exceed 1. Plot tick labels indicate the amino acid identities, with lysines (K) highlighted in bold. The percentage contribution of each carbohydrate type to the total glycanfibril hydrogen bonds is shown.

### 2 Torsional Configurational Entropy Analysis

#### 2.1 Workflow

The script `compute_entropy_N180.py` computes the torsional configurational entropy of an attached glycan using the Mutual Information Spanning Tree (MIST) formalism. It processes a multi-model PDB trajectory containing multiple conformations of the glycan and executes the following steps:

1. **Residue and connectivity definition.** The glycan residue range (`RESSEQ_RANGE`) and the inter-residue glycosidic linkages (`CONNECTIVITY_TEXT`) are specified to define the molecular graph.

2. **Structure parsing and atom selection.** Heavy atoms corresponding to the selected glycan residues are extracted from the topology using the `mdtraj` library.

3. **Bond and dihedral enumeration.** Intra-residue ring bonds and exocyclic heavy-atom bonds are identified automatically. User-defined inter-residue linkages are then incorporated to build the full heavy-atom connectivity graph, from which all rotatable dihedral angles (excluding ring bonds) are enumerated.

4. **Angle extraction and entropy estimation.** Dihedral time series are computed for each frame of the trajectory. One-dimensional torsional entropies and pairwise mutual informations are calculated from the binned angular distributions.

5. **MIST correction and total entropy.** A maximum-information spanning tree (MIST) is constructed from the mutual information matrix. The total torsional entropy is then estimated as:

$$S = \sum_i H_i - \sum_{(i,j)\in \mathrm{MST}} I_{ij}$$

in natural logarithmic units (*nats*) and in $S/k_B$.

6. **Output.** For each glycan segment analyzed, the script produces:

   - A CSV file listing all dihedral definitions and their individual entropies;
   - A CSV file of MIST edges and corresponding mutual information values;
   - A JSON summary file containing global entropy values and computation parameters;
   - A printed estimation of the free energy contribution at 300 K, computed as:

$$\Delta G_{300\,\mathrm{K}} = -0.593 \times (\Delta S/k_B)$$

   in $\mathrm{kcal\,mol^{-1}}$.

The main execution block automates the analysis over multiple contiguous glycan selections within the trajectory and reports the mean and standard deviation of the resulting entropic free-energy contributions.

The script was executed on 300 frames sampled with an even stride from the 1.5 $\mu$s MD simulations as:

```
python compute_entropy_N180.py full_traj_every_5.pdb > log_entropy_N180
```

#### 2.2 Results

Entropy calculation log for RML (N180 glycan, sialylated).

```
Processing selection 1: resSeq 1765 to 1782
  S/kB     : 40.664188
  dG_300K  : -24.113864 kcal/mol

Processing selection 2: resSeq 1747 to 1764
  S/kB     : 45.629522
  dG_300K  : -27.058307 kcal/mol

Processing selection 3: resSeq 1729 to 1746
  S/kB     : 38.170318
  dG_300K  : -22.634999 kcal/mol

Processing selection 4: resSeq 1711 to 1728
  S/kB     : 41.477473
  dG_300K  : -24.596142 kcal/mol

Processing selection 5: resSeq 1693 to 1710
  S/kB     : 43.751144
  dG_300K  : -25.944429 kcal/mol

Processing selection 6: resSeq 1675 to 1692
  S/kB     : 39.910851
  dG_300K  : -23.667135 kcal/mol

Processing selection 7: resSeq 1657 to 1674
  S/kB     : 38.950095
  dG_300K  : -23.097406 kcal/mol

Processing selection 8: resSeq 1639 to 1656
  S/kB     : 41.649705
  dG_300K  : -24.698275 kcal/mol

Processing selection 9: resSeq 1621 to 1638
  S/kB     : 40.017418
  dG_300K  : -23.730329 kcal/mol

Processing selection 10: resSeq 1783 to 1800
```

```
  S/kB     : 43.301650
  dG_300K  : -25.677878 kcal/mol

Processing selection 11: resSeq 1603 to 1620
  S/kB     : 40.412641
  dG_300K  : -23.964696 kcal/mol

Processing selection 12: resSeq 1585 to 1602
  S/kB     : 37.080079
  dG_300K  : -21.988487 kcal/mol

===== Summary of All Selections =====
 Start    End       S/kB      dG_300K
  1765   1782  40.664188   -24.113864
  1747   1764  45.629522   -27.058307
  1729   1746  38.170318   -22.634999
  1711   1728  41.477473   -24.596142
  1693   1710  43.751144   -25.944429
  1675   1692  39.910851   -23.667135
  1657   1674  38.950095   -23.097406
  1639   1656  41.649705   -24.698275
  1621   1638  40.017418   -23.730329
  1783   1800  43.301650   -25.677878
  1603   1620  40.412641   -23.964696
  1585   1602  37.080079   -21.988487
-------------------------------------
  MEAN         40.917924   -24.264329
   STD          2.328315     1.380691
```

#### Entropy calculation log for ME7 (N180 glycan, sialylated).

```
Processing selection 1: resSeq 1633 to 1650
  S/kB     : 37.145345
  dG_300K  : -22.027189 kcal/mol

Processing selection 2: resSeq 1651 to 1668
  S/kB     : 31.024648
  dG_300K  : -18.397616 kcal/mol

Processing selection 3: resSeq 1669 to 1686
  S/kB     : 40.738903
  dG_300K  : -24.158170 kcal/mol

Processing selection 4: resSeq 1687 to 1704
  S/kB     : 33.239344
  dG_300K  : -19.710931 kcal/mol

Processing selection 5: resSeq 1705 to 1722
  S/kB     : 37.211445
  dG_300K  : -22.066387 kcal/mol

Processing selection 6: resSeq 1723 to 1740
  S/kB     : 40.594308
  dG_300K  : -24.072425 kcal/mol
```

```
Processing selection 7: resSeq 1741 to 1758
  S/kB     : 32.159039
  dG_300K  : -19.070310 kcal/mol

Processing selection 8: resSeq 1759 to 1776
  S/kB     : 36.487763
  dG_300K  : -21.637244 kcal/mol

Processing selection 9: resSeq 1777 to 1794
  S/kB     : 40.793277
  dG_300K  : -24.190413 kcal/mol

Processing selection 10: resSeq 1795 to 1812
  S/kB     : 40.683202
  dG_300K  : -24.125139 kcal/mol

Processing selection 11: resSeq 1813 to 1830
  S/kB     : 43.518782
  dG_300K  : -25.806638 kcal/mol

Processing selection 12: resSeq 1831 to 1848
  S/kB     : 36.242599
  dG_300K  : -21.491861 kcal/mol

===== Summary of All Selections =====
 Start    End       S/kB      dG_300K
  1633   1650  37.145345   -22.027189
  1651   1668  31.024648   -18.397616
  1669   1686  40.738903   -24.158170
  1687   1704  33.239344   -19.710931
  1705   1722  37.211445   -22.066387
  1723   1740  40.594308   -24.072425
  1741   1758  32.159039   -19.070310
  1759   1776  36.487763   -21.637244
  1777   1794  40.793277   -24.190413
  1795   1812  40.683202   -24.125139
  1813   1830  43.518782   -25.806638
  1831   1848  36.242599   -21.491861
-------------------------------------
  MEAN         37.486555   -22.229527
   STD          3.749347     2.223363
```

#### 2.3 Estimation of the optimal binning size

The optimal number of bins for this analysis (72) was determined with the following `find_optimal_bins.py` script, that estimates optimal one- and two-dimensional histogram bin counts for torsional entropy calculations based on the MIST (Mutual Information Spanning Tree) formalism. It analyzes a multi-model PDB trajectory containing glycan conformations and provides per-selection and global recommendations for the number of angular bins.

1. **Residue selection and connectivity.** The user specifies the glycan residue ranges and the inter-residue glycosidic linkages. Residue ordinals are defined relative to the first residue within each selection.

2. **Trajectory parsing and atom selection.** The trajectory is read using `mdtraj`. Heavy atoms from the selected residues are identified, excluding hydrogens.

3. **Bond and dihedral construction.** Intra-residue ring and exocyclic heavy-atom bonds are automatically detected, and user-defined inter-residue linkages are added to construct the heavy-atom connectivity graph. All rotatable dihedral angles are enumerated, excluding ring-centered bonds.

4. **Angular analysis.** Dihedral angles are computed across all trajectory frames. For each dihedral distribution, the circular standard deviation and interquartile range are used to estimate optimal one-dimensional bin widths following the Scott, Freedman-Diaconis, and square-root rules adapted for circular variables:

$$h_S = 3.5\,\sigma_\theta N^{-1/3}, \quad h_F = 2.0\,\mathrm{IQR}_\theta N^{-1/3},$$

   leading to approximate bin counts $b_S = 2\pi/h_S$, $b_F = 2\pi/h_F$, and $b_Q = \sqrt{N}$. The final 1D bin count is the median of these estimates, further constrained to maintain a target mean occupancy of $\sim$20 samples per bin.

5. **2D bin recommendation.** Two-dimensional bin counts (`nbins_2d`) are derived from the total number of samples, enforcing an average of $\sim$30 samples per 2D cell, with the condition that `nbins_2d` $\leq$ `nbins_1d`.

6. **Output.** For each glycan selection, the script reports:

   - the number of frames and torsional angles analyzed,

- recommended `nbins_1d` and `nbins_2d` values,
- an aggregated global recommendation across all selections.

All results are written to a JSON file (`_bins_summary.json`) containing detailed per-selection and global binning parameters.

This utility provides consistent and statistically justified binning parameters for use in MIST-based torsional entropy analyses, ensuring comparable entropy estimates across multiple glycan segments.

#### 2.4 Scripts

##### find_optimal_bins.py

```
#!/usr/bin/env python3
# -*- coding: utf-8 -*-
"""
Estimate optimal 1D/2D histogram bin counts for torsional entropy (MIST)
from a multi-model PDB, using user-defined glycan selection and connectivity.

- Selection: inclusive PDB resSeq range(s)
- Connectivity: lines like 'bond 1.O6    18.C1' where residue ordinals start
  at 1 for the FIRST residue within the selection, in topology order.

Outputs:
- Per-selection recommended nbins_1d and nbins_2d
- Global recommendation across selections
- JSON summary

Requires: mdtraj, numpy
"""

import re, json, sys
import numpy as np
import mdtraj as md

# ------------------------- USER INPUT -----------------------------------------
PDB_PATH = "/mnt/data/rep1_reordered_stride_50.pdb"   # can be overridden by argv[1]
# Selections (match your attached scripts style)
STARTING_RESIDUES = [1633,  1651,   1669, 1687,     1705,   1723,   1741,   1759,   1777,   1795,   1813,   1831]
SELECTION_LENGTH  = 18  # residues per selection
OUT_PREFIX        = "bins_reco"

# Inter-residue connectivity (ordinals are 1-based within each selection)
CONNECTIVITY_TEXT = """
bond 1.O6    18.C1
bond 1.O4    2.C1
```

```
bond 2.O4    3.C1
bond 3.O6    8.C1
bond 3.O3    4.C1
bond 8.O6    12.C1
bond 8.O2    9.C1
bond 12.O4   13.C1
bond 13.O3   14.C1
bond 14.O4   15.C1
bond 14.O3   17.C1
bond 15.O3   16.C2
bond 9.O4    10.C1
bond 10.O6   11.C2
bond 4.O2    5.C1
bond 5.O4    6.C1
bond 6.O6    7.C2
""".strip()

# Targets and clamps for binning
NB1_MIN, NB1_MAX = 12, 180
NB2_MIN, NB2_MAX = 12, 60
TARGET_1D_COUNTS_PER_BIN = 20.0   # average samples per (occupied) 1D bin
TARGET_2D_COUNTS_PER_CELL = 30.0  # average samples per 2D cell

# ------------------------- UTILITIES ------------------------------------------
def _to_index_set(idx_like):
    out = []
    def add(x):
        if isinstance(x, (int, np.integer)): out.append(int(x))
        elif isinstance(x, (list, tuple, set, np.ndarray)):
            for y in x: add(y)
        else:
            try: out.append(int(x))
            except Exception: pass
    add(idx_like)
    return set(out)

def load_traj_pdb(path):
    t = md.load_pdb(path)
    print("N. frames", t.n_frames)
    if t.n_frames < 1: raise ValueError("No frames in PDB.")
    return t

def select_glycan_heavy(top, start_resseq, end_resseq):
    sel = top.select(f"(resSeq {start_resseq} to {end_resseq}) and not element H")
    if sel.size == 0: raise ValueError("Selection empty; check resSeq range and elements.")
    return sel

def residues_in_selection_order(top, heavy_idx):
    heavy = _to_index_set(heavy_idx)
    res = []
    seen = set()
    for r in top.residues:
        if any(a.index in heavy for a in r.atoms):
            if r.index not in seen:
                seen.add(r.index); res.append(r)
    return res  # ordinals = 1..len(res)
```

```
def parse_connectivity(text, res_list):
    """Parse 'bond X.Y   U.V' to atom-index pairs (i,j)."""
    edges = []
    line_re = re.compile(r"^\s*bond\s+(\S+)\s+(\S+)\s*$", re.IGNORECASE)
    def tok_to_index(tok):
        parts = tok.split(".")
        if len(parts) != 2: raise ValueError(f"Bad token '{tok}' (need 'ord.atom')")
        rord = int(parts[0]); aname = parts[1].strip()
        if not (1 <= rord <= len(res_list)):
            raise ValueError(f"Residue ordinal {rord} out of range 1..{len(res_list)}")
        res = res_list[rord-1]
        idx = None
        for a in res.atoms:
            if a.name.strip() == aname: idx = a.index; break
        if idx is None:
            au = aname.upper()
            for a in res.atoms:
                if a.name.strip().upper() == au: idx = a.index; break
        if idx is None:
            raise ValueError(f"Atom '{aname}' not found in residue ordinal {rord} ({res.name} {res.resSeq})")
        return idx
    for line in text.splitlines():
        if not line.strip() or line.strip().startswith("#"): continue
        m = line_re.match(line.strip())
        if not m: raise ValueError(f"Unrecognized connectivity line: '{line}'")
        i, j = tok_to_index(m.group(1)), tok_to_index(m.group(2))
        if i != j: edges.append((min(i,j), max(i,j)))
    return edges

def ring_bonds_by_names(top, allowed_idx):
    """Six-member ring bonds per residue name."""
    allowed = _to_index_set(allowed_idx)
    ring_pairs_default = [("C1","O5"),("O5","C5"),("C5","C4"),("C4","C3"),("C3","C2"),("C2","C1")]
    ring_pairs_OSA     = [("C2","O6"),("O6","C6"),("C6","C5"),("C5","C4"),("C4","C3"),("C3","C2")]
    ring = set()
    res_ids = set(top.atom(i).residue.index for i in allowed)
    for ridx in res_ids:
        res = list(top.residues)[ridx]
        names = {a.name.strip(): a.index for a in res.atoms
                  if a.index in allowed and (a.element is None or a.element.symbol != "H")}
        pairs = ring_pairs_OSA if res.name.strip() == "OSA" else ring_pairs_default
        for n1, n2 in pairs:
            if n1 in names and n2 in names:
                i, j = names[n1], names[n2]
                ring.add(frozenset((i, j)))
    return ring

def exocyclic_heavy_bonds_by_names(top, allowed_idx):
    """Minimal intra-residue heavy bonds useful for torsions."""
    allowed = _to_index_set(allowed_idx)
    bonds = set()
    res_ids = set(top.atom(i).residue.index for i in allowed)
    for ridx in res_ids:
        res = list(top.residues)[ridx]
        nm  = res.name.strip()
```

```
        names = {a.name.strip(): a.index for a in res.atoms
                 if a.index in allowed and (a.element is None or a.element.symbol != "H")}
        pairs = [("C4","O4"),("C3","O3")]
        if nm == "OSA": pairs += [("C2","C1"),("C1","O1"),("C5","O5")]
        else:           pairs += [("C5","C6"),("C6","O6"),("C2","O2"),("C1","O1")]
        for n1, n2 in pairs:
            if n1 in names and n2 in names:
                i, j = names[n1], names[n2]
                bonds.add(frozenset((i, j)))
    return bonds

def build_adjacency(top, heavy_idx, connectivity_pairs):
    heavy = _to_index_set(heavy_idx)
    adj = {i: set() for i in heavy}
    ring_b = ring_bonds_by_names(top, heavy)
    exo_b  = exocyclic_heavy_bonds_by_names(top, heavy)
    def add(i,j):
        if i in adj and j in adj: adj[i].add(j); adj[j].add(i)
    for e in ring_b | exo_b:
        i, j = tuple(e); add(i,j)
    for (i,j) in connectivity_pairs:
        add(i,j)
    return adj, ring_b

def enumerate_dihedrals(adj, ring_bonds):
    """All (a,b,c,d) with b-c edge, not a ring central bond."""
    D = set()
    for b, nbrs in adj.items():
        for c in nbrs:
            if b >= c: continue
            if frozenset((b,c)) in ring_bonds: continue
            As = [x for x in adj[b] if x != c]
            Ds = [x for x in adj[c] if x != b]
            if not As or not Ds: continue
            for a in As:
                for d in Ds:
                    if a == d or a == b or d == c: continue
                    D.add((int(a), int(b), int(c), int(d)))
    return sorted(D)

# ------------------------ CIRCULAR STATS  BINS -----------------------------
def circ_std(theta):
    N = theta.size
    C = np.sum(np.cos(theta))/N
    S = np.sum(np.sin(theta))/N
    R = np.hypot(C, S)
    return float(np.sqrt(max(0.0, -2.0*np.log(max(R, 1e-12)))))

def circ_iqr(theta):
    mu = np.arctan2(np.sum(np.sin(theta)), np.sum(np.cos(theta)))
    x = (theta - mu + np.pi) % (2*np.pi) - np.pi
    q25, q75 = np.quantile(x, 0.25), np.quantile(x, 0.75)
    dq = q75 - q25
    if dq > np.pi: dq = 2*np.pi - dq
    return float(dq)
```

```
def recommend_nbins_1d_for_angle(theta):
    N = int(theta.size)
    if N < 50: return 24
    sc = circ_std(theta)
    iq = circ_iqr(theta)
    hS = max(1e-3, 3.5*sc*(N**(-1/3)))
    hF = max(1e-3, 2.0*iq*(N**(-1/3)))
    bS = int(np.floor(2*np.pi/hS))
    bF = int(np.floor(2*np.pi/hF))
    bQ = int(np.floor(np.sqrt(N)))
    nb = int(np.median([bS, bF, bQ]))
    # enforce average occupancy target
    nb_cap = max(NB1_MIN, min(NB1_MAX, int(np.floor(N / TARGET_1D_COUNTS_PER_BIN)) if N >= TARGET_1D_COUNTS_PER_BIN else NB1_MIN))
    nb = int(np.clip(nb, NB1_MIN, min(NB1_MAX, nb_cap)))
    return nb

def recommend_global_bins(angle_matrix):
    """
    angle_matrix: shape (N_frames, N_dihedrals)
    Returns: nb1_global, nb2_global
    """
    N, D = angle_matrix.shape
    if D == 0 or N < 2:
        return NB1_MIN, NB2_MIN
    nb1_list = [recommend_nbins_1d_for_angle(angle_matrix[:, i]) for i in range(D)]
    nb1 = int(np.median(nb1_list))
    # 2D per-axis bins from target occupancy; clamp and keep leq nb1
    nb2_from_occup = int(np.floor(np.sqrt(N / TARGET_2D_COUNTS_PER_CELL))) if N > TARGET_2D_COUNTS_PER_CELL else NB2_MIN
    nb2 = int(np.clip(nb2_from_occup, NB2_MIN, NB2_MAX))
    nb2 = min(nb2, nb1)
    return nb1, nb2

# ------------------------ MAIN PIPELINE -------------------------------------
def build_angles_for_selection(traj, start_resseq, end_resseq):
    top = traj.topology
    heavy_idx = select_glycan_heavy(top, start_resseq, end_resseq)
    res_list  = residues_in_selection_order(top, heavy_idx)
    conn_pairs = parse_connectivity(CONNECTIVITY_TEXT, res_list)
    adj, ring_b = build_adjacency(top, heavy_idx, conn_pairs)
    diheds = enumerate_dihedrals(adj, ring_b)
    if not diheds:
        raise RuntimeError(f"No torsions found in selection {start_resseq}-{end_resseq}. Check connectivity.")
    dih_idx = np.array(diheds, dtype=int)
    angles = md.compute_dihedrals(traj, dih_idx)  # (N_frames, N_dihedrals)
    return angles, diheds

def main():
    pdb_path = PDB_PATH if len(sys.argv) < 2 else sys.argv[1]
    traj = load_traj_pdb(pdb_path)

    selections = [(s, s + SELECTION_LENGTH - 1) for s in STARTING_RESIDUES]
    recos = []
    for k, (rs, re) in enumerate(selections, 1):
        try:
            A, diheds = build_angles_for_selection(traj, rs, re)
            nb1, nb2 = recommend_global_bins(A)
```

```
            print(f"[sel {k:02d} | {rs}-{re}] frames={A.shape[0]} diheds={A.shape[1]}    nbins_1d={nb1}  nbins_2d={nb2}")
            recos.append((rs, re, A.shape[0], A.shape[1], nb1, nb2))
        except Exception as e:
            print(f"[sel {k:02d} | {rs}-{re}] ERROR: {e}")

    # Aggregate recommendation across selections
    if recos:
        nb1_all = [r[4] for r in recos]
        nb2_all = [r[5] for r in recos]
        nb1_global = int(np.median(nb1_all))
        nb2_global = int(np.median(nb2_all))
        print("\nRecommended global bins (use the SAME for both systems):")
        print(f"nbins_1d = {nb1_global}")
        print(f"nbins_2d = {nb2_global}")

        summary = {
            "pdb_path": pdb_path,
            "target_1d_avg_counts": TARGET_1D_COUNTS_PER_BIN,
            "target_2d_avg_counts": TARGET_2D_COUNTS_PER_CELL,
            "per_selection": [
                {"start": r[0], "end": r[1], "frames": r[2], "n_dihedrals": r[3],
                 "nbins_1d": r[4], "nbins_2d": r[5]} for r in recos
            ],
            "global": {"nbins_1d": nb1_global, "nbins_2d": nb2_global}
        }
        with open(f"{OUT_PREFIX}_bins_summary.json", "w") as f:
            json.dump(summary, f, indent=2)
        print(f"\nWrote {OUT_PREFIX}_bins_summary.json")

if __name__ == "__main__":
    main()
```

#### compute_entropy_N180.py

```
#!/usr/bin/env python3
# -*- coding: utf-8 -*-
"""
Torsional configurational entropy with MIST for an attached glycan.

Inputs:
- Multi-model PDB trajectory
- CONNECTIVITY_TEXT: inter-residue bonds, e.g. 'bond 1.O6    18.C1'

Outputs:
- S (nats and S/kB), dihedral table CSV, MST CSV, summary JSON
- deltaG_300K [kcal/mol] = -0.593 * (deltaS/kB)

Requires: mdtraj, numpy
"""

import re, json
import numpy as np
import mdtraj as md
```

```
# ------------------------ USER INPUT ----------------------------------------
OUT_PREFIX = "glycan_torsion_entropy_rep1"

CONNECTIVITY_TEXT = """
bond 1.O6    18.C1
bond 1.O4    2.C1
bond 2.O4  3.C1
bond 3.O6  8.C1
bond 3.O3  4.C1
bond 8.O6  12.C1
bond 8.O2  9.C1
bond 12.O4 13.C1
bond 13.O3 14.C1
bond 14.O4 15.C1
bond 14.O3 17.C1
bond 15.O3 16.C2
bond 9.O4  10.C1
bond 10.O6  11.C2
bond 4.O2  5.C1
bond 5.O4  6.C1
bond 6.O6  7.C2
""".strip()

# ------------------------ UTILS ---------------------------------------------
def _to_index_set(idx_like):
    """Flatten any nested iterable of indices into a set of ints."""
    out = []
    def add(x):
        if isinstance(x, (int, np.integer)):
            out.append(int(x))
        elif isinstance(x, (list, tuple, set, np.ndarray)):
            for y in x:
                add(y)
        else:
            try:
                out.append(int(x))
            except Exception:
                pass
    add(idx_like)
    return set(out)

# ------------------------ SELECTION -----------------------------------------
def load_traj_pdb(pdb_path):
    t = md.load_pdb(pdb_path)
    if t.n_frames < 1:
        raise ValueError("No frames in PDB.")
    return t

def select_glycan_heavy(top, start_resseq, end_resseq):
    sel = top.select(f"(resSeq {start_resseq} to {end_resseq}) and not element H")
    if sel.size == 0:
        raise ValueError("Selection empty; check RESSEQ_RANGE and elements.")
    return sel

def residues_in_selection_order(top, heavy_idx):
```

```
    heavy = _to_index_set(heavy_idx)
    res_list = []
    seen = set()
    for r in top.residues:
        if any(a.index in heavy for a in r.atoms):
            if r.index not in seen:
                seen.add(r.index)
                res_list.append(r)
    return res_list  # ordinals 1..N in topology order

# ------------------------ CONNECTIVITY --------------------------------------
def parse_connectivity(text, res_list):
    """
    Parse 'bond X.Y   U.V' where X,U are residue ordinals (1-based within res_list),
    and Y,V are atom names within those residues.
    Returns list of atom-index pairs (i,j).
    """
    edges = []
    line_re = re.compile(r"^\s*bond\s+(\S+)\s+(\S+)\s*$", re.IGNORECASE)

    def parse_token(tok):
        # token like '1.O6' or '18.C1'
        parts = tok.split(".")
        if len(parts) != 2:
            raise ValueError(f"Bad token: '{tok}' (expected 'ord.atom')")
        res_ord = int(parts[0])
        atom_name = parts[1].strip()
        if not (1 <= res_ord <= len(res_list)):
            raise ValueError(f"Residue ordinal {res_ord} out of range [1..{len(res_list)}]")
        res = res_list[res_ord - 1]
        # exact name match, then case-insensitive fallback
        idx = None
        for a in res.atoms:
            if a.name.strip() == atom_name:
                idx = a.index; break
        if idx is None:
            atom_name_u = atom_name.upper()
            for a in res.atoms:
                if a.name.strip().upper() == atom_name_u:
                    idx = a.index; break
        if idx is None:
            raise ValueError(f"Atom '{atom_name}' not found in residue ordinal {res_ord} ({res.name} {res.resSeq})")
        return idx

    for line in text.splitlines():
        line = line.strip()
        if not line or line.startswith("#"):
            continue
        m = line_re.match(line)
        if not m:
            raise ValueError(f"Unrecognized connectivity line: '{line}'")
        t1, t2 = m.group(1), m.group(2)
        i = parse_token(t1)
        j = parse_token(t2)
        if i != j:
            edges.append((min(i, j), max(i, j)))
```

```python
    return edges

# ------------------------ INTRA-RESIDUE BONDS --------------------------------
def ring_bonds_by_names(top, allowed_idx):
    """
    Intra-residue ring bonds by atom names.
    Residue != 'OSA': ring atoms {C1,C2,C3,C4,C5,O5} with bonds:
        C1-O5, O5-C5, C5-C4, C4-C3, C3-C2, C2-C1
    Residue == 'OSA': ring atoms {C2,C3,C4,C5,C6,O6} with bonds:
        C2-O6, O6-C6, C6-C5, C5-C4, C4-C3, C3-C2
    Returns set of frozenset({i,j}) restricted to allowed_idx.
    """
    allowed = _to_index_set(allowed_idx)
    ring_pairs_default = [("C1","O5"),("O5","C5"),("C5","C4"),("C4","C3"),("C3","C2"),("C2","C1")]
    ring_pairs_OSA     = [("C2","O6"),("O6","C6"),("C6","C5"),("C5","C4"),("C4","C3"),("C3","C2")]
    ring_bonds = set()
    res_ids = set(md.Topology.atom(top, i).residue.index for i in allowed)
    for ridx in res_ids:
        res = list(top.residues)[ridx]
        names_to_idx = {a.name.strip(): a.index
                        for a in res.atoms
                        if a.index in allowed and (a.element is None or a.element.symbol != "H")}
        pairs = ring_pairs_OSA if res.name.strip() == "OSA" else ring_pairs_default
        for n1, n2 in pairs:
            if n1 in names_to_idx and n2 in names_to_idx:
                i, j = names_to_idx[n1], names_to_idx[n2]
                ring_bonds.add(frozenset((i, j)))
    return ring_bonds

def exocyclic_heavy_bonds_by_names(top, allowed_idx):
    """
    Minimal intra-residue heavy bonds for torsions.
    Default: C5-C6, C6-O6, C4-O4, C3-O3, C2-O2, C1-O1 (if present)
    OSA   : C2-C1, C1-O1, C4-O4, C3-O3, C5-O5 (if present)
    """
    allowed = _to_index_set(allowed_idx)
    bonds = set()
    res_ids = set(md.Topology.atom(top, i).residue.index for i in allowed)
    for ridx in res_ids:
        res = list(top.residues)[ridx]
        nm = res.name.strip()
        names_to_idx = {a.name.strip(): a.index
                        for a in res.atoms
                        if a.index in allowed and (a.element is None or a.element.symbol != "H")}
        pairs = [("C4","O4"),("C3","O3")]  # common subset
        if nm == "OSA":
            pairs += [("C2","C1"),("C1","O1"),("C5","O5")]
        else:
            pairs += [("C5","C6"),("C6","O6"),("C2","O2"),("C1","O1")]
        for n1, n2 in pairs:
            if n1 in names_to_idx and n2 in names_to_idx:
                i, j = names_to_idx[n1], names_to_idx[n2]
                bonds.add(frozenset((i, j)))
    return bonds

# ------------------------ GRAPH + DIHEDRALS ---------------------------------
```

```
def build_adjacency(top, heavy_idx, connectivity_pairs):
    """
    Undirected adjacency among heavy atoms:
      - ring bonds (for neighbors, but excluded as central bonds),
      - minimal exocyclic heavy bonds,
      - user-specified inter-residue linkages.
    """
    heavy = _to_index_set(heavy_idx)
    adj = {i: set() for i in heavy}

    ring_b = ring_bonds_by_names(top, heavy)
    exo_b  = exocyclic_heavy_bonds_by_names(top, heavy)

    def add_edge(i, j):
        if i in adj and j in adj:
            adj[i].add(j); adj[j].add(i)

    for e in ring_b | exo_b:
        i, j = tuple(e)
        add_edge(i, j)

    for (i, j) in connectivity_pairs:
        add_edge(i, j)

    return adj, ring_b

def enumerate_dihedrals(adj, ring_bonds):
    """All a-b-c-d with b-c in adj, excluding ring bonds as central."""
    dihed = set()
    for b, nbrs in adj.items():
        for c in nbrs:
            if b >= c:
                continue
            if frozenset((b, c)) in ring_bonds:
                continue
            As = [x for x in adj[b] if x != c]
            Ds = [x for x in adj[c] if x != b]
            if not As or not Ds:
                continue
            for a in As:
                for d in Ds:
                    if a == d or a == b or d == c:
                        continue
                    dihed.add((int(a), int(b), int(c), int(d)))
    return sorted(dihed)

# ------------------------ ENTROPY + MIST ------------------------------------
def entropy_1d_circular(angles, nbins=72):
    N = int(angles.size)
    if N < 2:
        return 0.0
    counts, _ = np.histogram(angles, bins=nbins, range=(-np.pi, np.pi))
    nz = counts[counts > 0]
    if nz.size == 0:
        return 0.0
    p = nz.astype(np.float64) / float(N)
```

```
    H = -np.sum(p * np.log(p))
    K = int(nz.size)
    H += (K - 1) / (2.0 * N)  # MillerMadow
    return float(H)

def entropy_2d_torus(a1, a2, nbins=36):
    if a1.size != a2.size:
        raise ValueError("Angle arrays must have same length.")
    N = int(a1.size)
    if N < 2:
        return 0.0
    counts, _, _ = np.histogram2d(a1, a2, bins=nbins,
                                  range=[(-np.pi, np.pi), (-np.pi, np.pi)])
    nz = counts[counts > 0]
    if nz.size == 0:
        return 0.0
    p = nz.astype(np.float64) / float(N)
    H = -np.sum(p * np.log(p))
    K = int(nz.size)
    H += (K - 1) / (2.0 * N)
    return float(H)

def mutual_information_torus(a1, a2, nbins_1d=72, nbins_2d=36):
    H1 = entropy_1d_circular(a1, nbins_1d)
    H2 = entropy_1d_circular(a2, nbins_1d)
    H12 = entropy_2d_torus(a1, a2, nbins_2d)
    I = H1 + H2 - H12
    return float(max(0.0, I))

def maximum_information_spanning_tree(I):
    n = int(I.shape[0])
    if n == 0:
        return []
    sel = np.zeros(n, dtype=bool); sel[0] = True
    edges = []
    for _ in range(n - 1):
        si = np.where(sel)[0]; ui = np.where(~sel)[0]
        if ui.size == 0:
            break
        best_w, best_u, best_v = -1.0, -1, -1
        for u in si:
            row = I[u, ui]
            j = int(np.argmax(row))
            w = float(row[j])
            if w > best_w:
                best_w, best_u, best_v = w, u, int(ui[j])
        if best_u < 0:
            best_u, best_v = int(si[0]), int(ui[0])
        sel[best_v] = True
        edges.append((min(best_u, best_v), max(best_u, best_v)))
    return edges

# ------------------------- PIPELINE -------------------------------------------
def torsion_entropy_MIST(pdb_path, resseq_range, out_prefix,
                         nbins_1d=72, nbins_2d=36, connectivity_text=""):
    traj = load_traj_pdb(pdb_path)
```

```
    top = traj.topology
    heavy_idx = select_glycan_heavy(top, resseq_range[0], resseq_range[1])
    res_list = residues_in_selection_order(top, heavy_idx)

    # Build adjacency graph
    conn_pairs = parse_connectivity(connectivity_text, res_list)
    adj, ring_b = build_adjacency(top, heavy_idx, conn_pairs)

    # Dihedrals and angles
    diheds = enumerate_dihedrals(adj, ring_b)
    if not diheds:
        raise RuntimeError("No rotatable heavy-atom dihedrals from provided connectivity.")
    dih_idx = np.array(diheds, dtype=int)
    angles = md.compute_dihedrals(traj, dih_idx)  # (n_frames, n_dihedrals)

    nF, nD = angles.shape
    H = np.array([entropy_1d_circular(angles[:, i], nbins_1d) for i in range(nD)], dtype=float)

    I = np.zeros((nD, nD), dtype=float)
    for i in range(nD):
        ai = angles[:, i]
        for j in range(i + 1, nD):
            Iij = mutual_information_torus(ai, angles[:, j], nbins_1d, nbins_2d)
            I[i, j] = Iij; I[j, i] = Iij

    mst_edges = maximum_information_spanning_tree(I)
    sum_H = float(np.sum(H))
    sum_I_tree = float(np.sum([I[u, v] for (u, v) in mst_edges])) if mst_edges else 0.0
    S_nats = sum_H - sum_I_tree
    S_over_kB = S_nats  # natural logs  numeric equality

    # Outputs
    rows = []
    for k, (a, b, c, d) in enumerate(diheds):
        aa, bb, cc, dd = top.atom(a), top.atom(b), top.atom(c), top.atom(d)
        rows.append([
            k, a, aa.name, aa.residue.name, aa.residue.resSeq,
            b, bb.name, bb.residue.name, bb.residue.resSeq,
            c, cc.name, cc.residue.name, cc.residue.resSeq,
            d, dd.name, dd.residue.name, dd.residue.resSeq,
            H[k]
        ])
    header = ["idx",
              "a_idx","a_name","a_resn","a_resSeq",
              "b_idx","b_name","b_resn","b_resSeq",
              "c_idx","c_name","c_resn","c_resSeq",
              "d_idx","d_name","d_resn","d_resSeq",
              "H_i_nats"]
#    np.savetxt(f"{out_prefix}_dihedrals.csv", np.array(rows, dtype=object),
#               fmt="%s", delimiter=",", header=",".join(header), comments="")

    mst_rows = [[u, v, I[u, v]] for (u, v) in mst_edges]
#    np.savetxt(f"{out_prefix}_MST_edges.csv", np.array(mst_rows, dtype=object),
#               fmt="%s", delimiter=",", header="u_idx,v_idx,I_uv_nats", comments="")

    summary = dict(
```

```
        pdb_path=pdb_path,
        resseq_start=int(resseq_range[0]),
        resseq_end=int(resseq_range[1]),
        n_frames=int(nF),
        n_dihedrals=int(nD),
        nbins_1d=int(nbins_1d),
        nbins_2d=int(nbins_2d),
        S_nats=float(S_nats),
        S_over_kB=float(S_over_kB),
        kBT_300K_kcal=0.593,
        note="Compute per system. deltaG_300K [kcal/mol] = -0.593 * (deltaS/kB)."
    )
    with open(f"{out_prefix}_summary.json", "w") as f:
        json.dump(summary, f, indent=2)

    return dict(S_nats=S_nats, S_over_kB=S_over_kB,
                N_frames=nF, N_dihedrals=nD,
                dihedrals=diheds, H=H, I=I, mst_edges=mst_edges)

# ------------------------ MAIN ---------------------------------------------
if __name__ == "__main__":
    import sys

    if len(sys.argv) < 2:
        print("Usage: python compute_entropy.py <PDB_PATH>")
        sys.exit(1)

    PDB_PATH = sys.argv[1]

    # Example configuration: adjust these as needed
    STARTING_RESIDUES = [1765,1747,1729,1711,1693,1675,1657,1639,1621,1783,1603,1585]
    SELECTION_LENGTH = 18  # number of residues per selection
    OUT_PREFIX = "glycan_entropy_N180"

    all_results = []

    for i, start_resseq in enumerate(STARTING_RESIDUES):
        end_resseq = start_resseq + SELECTION_LENGTH - 1
        out_prefix_i = f"{OUT_PREFIX}_sel{i+1}_{start_resseq}_{end_resseq}"

        print(f"\nProcessing selection {i+1}: resSeq {start_resseq} to {end_resseq}")
        try:
            result = torsion_entropy_MIST(
                pdb_path=PDB_PATH,
                resseq_range=(start_resseq, end_resseq),
                out_prefix=out_prefix_i,
                nbins_1d=15,
                nbins_2d=12,
                connectivity_text=CONNECTIVITY_TEXT
            )
            S = result['S_over_kB']
            deltaG = -0.593 * S
            all_results.append((start_resseq, end_resseq, S, deltaG))
            print(f"  S/kB     : {S:.6f}")
            print(f"  deltaG_300K  : {deltaG:.6f} kcal/mol")
        except Exception as e:
```

```
            print(f"  Error in selection {start_resseq}-{end_resseq}: {e}")

    if all_results:
        Ss = [r[2] for r in all_results]
        Gs = [r[3] for r in all_results]
        mean_S = np.mean(Ss)
        std_S = np.std(Ss)
        mean_G = np.mean(Gs)
        std_G = np.std(Gs)

        print("\n===== Summary of All Selections =====")
        print(f"{'Start':>6} {'End':>6} {'S/kB':>10} {'deltaG_300K':>12}")
        for start, end, S, G in all_results:
            print(f"{start:6} {end:6} {S:10.6f} {G:12.6f}")
        print("------------------------------------")
        print(f"{'MEAN':>6} {'':6} {mean_S:10.6f} {mean_G:12.6f}")
        print(f"{'STD':>6} {'':6} {std_S:10.6f} {std_G:12.6f}")
```

### 3 Sample NCIPLOT4 input

NCIPLOT calculations were run on 30 frames sampled with an even stride from the 1.5 $\mu$s MD simulations of RML and ME7 fibrils glycosylated with sialylated glycans. For each frame, we ran an independent NCIPLOT calculation and plotted all $s$ vs $sign(\lambda_2)\rho$ collectively (plots in Figure 4 of the main text). A sample input is shown below.

```
2
frag1.xyz
frag2.xyz
CG2FG 4 8 4 2 1
INCREMENTS 0.5 0.5 0.5
INTERMOLECULAR
OUTPUT 1
```

### 4 Sample script for strand reordering

Owing to diffusion, wrapping of fibril coordinates into the primary simulation box (`iwrap=1` in Amber) can lead to a change of the ordering of the strands along the simulation. Prior to analysis, strands are reordered using the following script.

```
import glob
from math import sqrt
from Bio.PDB import PDBParser, PDBIO, Chain, Atom
import os
import sys
from shutil import move, rmtree
import natsort

def split_pdb_into_frames(input_pdb, output_directory):
    if not os.path.exists(output_directory):
        os.makedirs(output_directory)

    with open(input_pdb, 'r') as pdb_file:
        frame_lines = []
        frame_count = 0

        for line in pdb_file:
            if line.startswith('MODEL'):
                frame_lines = [line]
                frame_count += 1
            elif line.startswith('ENDMDL'):
                frame_lines.append(line)
                output_file = os.path.join(output_directory, f'frame_{frame_count}.pdb')
                with open(output_file, 'w') as frame_file:
                    frame_file.writelines(frame_lines)
                frame_lines = []
            else:
                if frame_lines:
                    frame_lines.append(line)

    print(f"Split {frame_count} frames into individual PDB files in {output_directory}.")

def find_seventh_ca_line_index(pdb_content):
    ca_count = 0
    for index, line in enumerate(pdb_content.splitlines()):
        if line.startswith("ATOM") and " CA " in line[12:16]:
            ca_count += 1
            if ca_count == 7:
                return index
    return None
def extract_chains_with_glycans_v2(frame_pdb):
    protein_chains = []
    glycans = []
    chain_lines = []
    glycan_lines = []
    is_glycan = False
    h2o_ofa_counter = 0  # Counter to track "H2O 0fA" appearances

    with open(frame_pdb, 'r') as pdb_file:
        for line in pdb_file:
            if ' OXT ' in line:  # End of a protein chain
                chain_lines.append(line)
                protein_chains.append("".join(chain_lines))
                chain_lines = []
```

```
        elif ' UYB ' in line and ' C1 ' in line:  # Start of a glycan
            is_glycan = True
            glycan_lines = [line]
            h2o_ofa_counter = 0  # Reset counter
        elif is_glycan:
            glycan_lines.append(line)

            # Detect "H2O OfA" and count occurrences
            if ' H2O ' in line and ' OfA ' in line:
                h2o_ofa_counter += 1

            # End glycan when "H2O OfA" appears twice
            if h2o_ofa_counter == 2:
                glycans.append("".join(glycan_lines))
                glycan_lines = []
                is_glycan = False
        else:
            chain_lines.append(line)

if glycan_lines:  # Ensure last glycan is saved
    glycans.append("".join(glycan_lines))

# Associate glycans with protein chains
combined_chains = []
for protein_chain in protein_chains:
    nln_lines = [line for line in protein_chain.splitlines() if ' NLN ' in line and ' ND2 ' in line]
    num_glycans_needed = len(nln_lines)

    if num_glycans_needed == 0:
        combined_chains.append(protein_chain)
        continue

    nd2_coords_list = [list(map(float, line[30:54].split())) for line in nln_lines]

    assigned_glycans = []
    used_glycans = set()

    for nd2_coords in nd2_coords_list:
        min_distance = float('inf')
        best_glycan = None

        for i, glycan in enumerate(glycans):
            if i in used_glycans:
                continue

            c1_lines = [line for line in glycan.splitlines() if ' UYB ' in line and ' C1 ' in line]
            for c1_line in c1_lines:
                c1_coords = list(map(float, c1_line[30:54].split()))
                distance = sqrt(sum((a - b) ** 2 for a, b in zip(nd2_coords, c1_coords)))

                if distance < min_distance:
                    min_distance = distance
                    best_glycan = glycan
                    best_glycan_index = i

        if best_glycan:
```

```
                assigned_glycans.append(best_glycan)
                used_glycans.add(best_glycan_index)

        combined_chains.append(protein_chain + "\n" + "\n".join(assigned_glycans))

    return combined_chains

def trim_chains_to_atom_info(chains):
    return ["\n".join(line[:30] for line in chain.splitlines() if line.startswith("ATOM")) for chain in chains]

def trim_chains_to_coordinates(chains):
    return ["\n".join(line[30:] for line in chain.splitlines() if line.startswith("ATOM")) for chain in chains]

def reorder_trimmed_coordinates_by_z(trimmed_coordinates, line_index):
    def extract_z_coordinate(chain):
        try:
            lines = chain.splitlines()
            return float(lines[line_index].split()[2])
        except (IndexError, ValueError):
            return float('inf')

    indexed_coords = list(enumerate(trimmed_coordinates))
    indexed_coords.sort(key=lambda x: extract_z_coordinate(x[1]))

    sorted_coordinates = [coord for _, coord in indexed_coords]
    order_mapping = [index for index, _ in indexed_coords]

    return sorted_coordinates, order_mapping

def combine_atom_info_and_coordinates(atom_info, reordered_coordinates, order_mapping):
    if len(atom_info) != len(reordered_coordinates):
        raise ValueError("Atom info and reordered coordinates must have the same number of elements.")

    reordered_atom_info = [atom_info[i] for i in order_mapping]

    combined = []
    for atom_block, coord_block in zip(reordered_atom_info, reordered_coordinates):
        atom_lines = atom_block.splitlines()
        coord_lines = coord_block.splitlines()

        if len(atom_lines) != len(coord_lines):
            raise ValueError("Mismatch in the number of lines between atom info and coordinates for a chain.")

        combined_block = "\n".join(f"{atom}{coord}" for atom, coord in zip(atom_lines, coord_lines))
        combined.append(combined_block)

    return combined

def merge_files_with_custom_joiner(file_names, output_file, joiner):
    with open(output_file, 'w') as out_file:
        for i, file_name in enumerate(file_names):
            with open(file_name, 'r') as in_file:
                content = in_file.read()
                out_file.write(content)
                if i < len(file_names) - 1:
```

```
                out_file.write(joiner)

    print(f"Merged {len(file_names)} files into {output_file}.")

pdb_file = sys.argv[1]
stride = int(sys.argv[2])

split_pdb_into_frames(pdb_file, "pieces")
frames = natsort.natsorted(glob.glob("pieces/*pdb"))

written_frames = []

for frame_number, frame in enumerate(frames, start=1):
    if "reordered" not in frame:
        combined_chains = extract_chains_with_glycans_v2(frame)
        atom_info = trim_chains_to_atom_info(combined_chains)
        coords = trim_chains_to_coordinates(combined_chains)

        if combined_chains:
            index_for_reorder = find_seventh_ca_line_index(combined_chains[0])
            if index_for_reorder is not None:
                reordered_coords, order_mapping = reorder_trimmed_coordinates_by_z(coords, index_for_reorder)
                reordered_contents = combine_atom_info_and_coordinates(atom_info, reordered_coords, order_mapping)
            else:
                reordered_contents = combine_atom_info_and_coordinates(atom_info, coords, list(range(len(coords))))
        else:
            reordered_contents = []

        if frame_number % stride == 0:
            output_frame = frame.replace("frame", "reordered_frame")
            written_frames.append(output_frame)

            with open(output_frame, "w") as g:
                g.write(f"MODEL {frame_number}\n")
                g.write("\nTER\n".join(reordered_contents))
                g.write("\nENDMDL\n")

to_merge = natsort.natsorted(written_frames)
merge_files_with_custom_joiner(to_merge, pdb_file.replace(".pdb", f"_reordered_stride_{stride}.pdb"), "")
rmtree("pieces")
```

### 5 Sample script for C$\alpha$ RMSF calculation

The following script was used to compute per-residue C$\alpha$ Root-Mean-Square Fluctuation (RMSF) values.

```
import numpy as np
import sys
import re
```

```
from collections import defaultdict

def compute_alpha_carbon_rmsf(pdb_file, output_file):
    # Initialize data structures
    residue_positions = defaultdict(list)
    residue_numbers = set()

    with open(pdb_file, 'r') as file:
        current_frame = []
        for line in file:
            if line.startswith("MODEL"):
                current_frame = []  # Start a new frame
            elif line.startswith("ATOM") and line[13:15] == "CA":
#                print(line)
                # Extract residue number and coordinates
                residue_number = int(line[22:26].strip())
                x = float(line[30:38].strip())
                y = float(line[38:46].strip())
                z = float(line[46:54].strip())
                residue_positions[residue_number].append(np.array([x, y, z]))
                residue_numbers.add(residue_number)
            elif line.startswith("ENDMDL"):
                # End of the current frame
                pass

    # Sort residue numbers for consistent output
    #residue_numbers = sorted(residue_numbers)

    # Compute RMSF
    rmsf_values = {}
    for residue_number in residue_numbers:
        positions = np.array(residue_positions[residue_number])
        mean_position = np.mean(positions, axis=0)
        squared_displacements = np.sum((positions - mean_position) ** 2, axis=1)
        rmsf_values[residue_number] = np.sqrt(np.mean(squared_displacements))

    # Write RMSF to output file
    with open(output_file, 'w') as out_file:
        for residue_number in residue_numbers:
            out_file.write(f"{residue_number:.3f}\t{rmsf_values[residue_number]:.4f}\n")

    print(f"RMSF values have been saved to {output_file}.")

# Example usage
pdb_file = sys.argv[1]  # Replace with your PDB file path
output_file = pdb_file.replace(".pdb", "")+"_alpha_carbon_rmsf.txt"  # Output file
compute_alpha_carbon_rmsf(pdb_file, output_file)
```

### 6 Sample script for glycans RMSF calculation

The following script was used to compute per-carbohydrate unit C1 RMSF values.

```
import numpy as np
```

```
import sys
import re
from collections import defaultdict

def compute_alpha_carbon_rmsf(pdb_file, output_file):
    # Initialize data structures
    residue_positions = defaultdict(list)
    residue_numbers = set()

    with open(pdb_file, 'r') as file:
        current_frame = []
        for line in file:
            if line.startswith("MODEL"):
                current_frame = []  # Start a new frame
            elif line.startswith("ATOM") and line.strip()[17:20] \
              ['0fA', 'VMB', '4YB', '0SA', 'WYB', '3LB', 'XMA', '6LB', 'UYB', '2MA'] and line.strip()[12:16].strip() == "C1":
#                print(line)
                # Extract residue number and coordinates
                residue_number = int(line[22:26].strip())
                x = float(line[30:38].strip())
                y = float(line[38:46].strip())
                z = float(line[46:54].strip())
                residue_positions[residue_number].append(np.array([x, y, z]))
                residue_numbers.add(residue_number)
            elif line.startswith("ENDMDL"):
                # End of the current frame
                pass

    # Sort residue numbers for consistent output
    #residue_numbers = sorted(residue_numbers)

    # Compute RMSF
    rmsf_values = {}
    for residue_number in residue_numbers:
        positions = np.array(residue_positions[residue_number])
        mean_position = np.mean(positions, axis=0)
        squared_displacements = np.sum((positions - mean_position) ** 2, axis=1)
        rmsf_values[residue_number] = np.sqrt(np.mean(squared_displacements))

    # Write RMSF to output file
    with open(output_file, 'w') as out_file:
        for residue_number in residue_numbers:
            out_file.write(f"{residue_number:.3f}\t{rmsf_values[residue_number]:.4f}\n")

    print(f"RMSF values have been saved to {output_file}.")

# Example usage
pdb_file = sys.argv[1]  # Replace with your PDB file path
output_file = pdb_file.replace(".pdb", "")+"_carbos_rmsf.txt"  # Output file
compute_alpha_carbon_rmsf(pdb_file, output_file)
```

### 7 Sample script for per-residue SASA analysis

The following script was used to compute per-residue Solvent Accessible Surface Area (SASA) values.

```
#!/usr/bin/env python3
import argparse, os, tempfile
from collections import defaultdict
from typing import List, Dict
import pandas as pd
import freesasa

def split_models(pdb_path: str) -> List[str]:
    models, current = [], []
    with open(pdb_path) as f:
        for line in f:
            rec = line[0:6].strip().upper()
            if rec == "MODEL":
                if current:
                    models.append("".join(current))
                    current = []
            elif rec == "ENDMDL":
                if current:
                    models.append("".join(current))
                    current = []
            else:
                current.append(line)
    if current:
        models.append("".join(current))
    if not models:
        with open(pdb_path) as f:
            models = ["".join(f.readlines())]
    return models

def sanitize_pdb_block(pdb_text: str) -> str:
    """Keep only properly formatted ATOM/HETATM lines"""
    lines = []
    for l in pdb_text.splitlines():
        if l[0:6].strip().upper() in ("ATOM", "HETATM"):
            lines.append(l)
    return "\n".join(lines) + "\n" if lines else ""

def sasa_per_residue_for_frame(pdb_text: str) -> List[Dict]:
    pdb_text = sanitize_pdb_block(pdb_text)
    if not pdb_text.strip():
        return []  # empty model, skip

    with tempfile.NamedTemporaryFile(delete=False, suffix=".pdb", mode="w") as tmp:
        tmp.write(pdb_text)
        tmp_name = tmp.name

    try:
        structure = freesasa.Structure(tmp_name)
```

```
        result = freesasa.calc(structure)
    finally:
        try:
            os.remove(tmp_name)
        except OSError:
            pass

    per_res = defaultdict(float)
    for i in range(structure.nAtoms()):
        per_res[
            (
                structure.chainLabel(i).strip(),
                structure.residueNumber(i).strip(),
                structure.residueName(i).strip(),
            )
        ] += float(result.atomArea(i))

    return [
        {
            "Chain": c,
            "Residue_ID": r,
            "Residue_Name": rn,
            "SASA(^2)": area,
        }
        for (c, r, rn), area in per_res.items()
    ]

def main():
    ap = argparse.ArgumentParser()
    ap.add_argument("-i", "--pdb", required=True)
    ap.add_argument("-o", "--out-per-frame", default="sasa_per_residue.csv")
    ap.add_argument("-a", "--out-average", default="sasa_per_residue_average.csv")
    args = ap.parse_args()

    models = split_models(args.pdb)
    all_rows = []

    for idx, m in enumerate(models, 1):
        rows = sasa_per_residue_for_frame(m)
        if not rows:
            print(f"Skipping empty frame {idx}")
            continue
        for r in rows:
            r["Frame"] = idx
        all_rows.extend(rows)

    if not all_rows:
        raise RuntimeError("No valid frames contained ATOM/HETATM records")

    df = pd.DataFrame(all_rows)
    df.to_csv(args.out_per_frame, index=False)

    df_mean = (
        df.groupby(["Chain", "Residue_ID", "Residue_Name"], as_index=False)["SASA(^2)"]
        .mean()
```

```
        .rename(columns={"SASA(^2)": "Mean_SASA(^2)"})
    )
    df_mean.to_csv(args.out_average, index=False)

if __name__ == "__main__":
    main()
```

### 8 Sample script for inter-strand Cα distances

Cα distances between equivalent residues on consecutive strands of the fibril were computed using the following script.

```
 #!/usr/bin/env python3
"""
Compute interchain CC distances for each MODEL of a multimodel PDB.

Modifications
-------------
* Robust chain splitting: every line whose record name (columns 16) is TER
  triggers a new chain. Duplicate TER lines never generate empty chains.
* Stricter validation and exhaustively documented, PEP484typed code.
"""

from __future__ import annotations

import sys
from pathlib import Path
from typing import List

import numpy as np

BUFFER_RES: int = 0          # residues to ignore at each terminus (flexible ends)

def read_pdb_frames(pdb_path: str) -> List[str]:
    """Return each MODELENDMDL block as a raw string."""
    frames: List[str] = []
    current: list[str] = []

    with Path(pdb_path).open() as fh:
        for line in fh:
            if line.startswith("MODEL"):
                current = []
            current.append(line.rstrip("\n"))
            if line.startswith("ENDMDL"):
                frames.append("\n".join(current))

    return frames
```

```
def split_chains(frame: str) -> List[str]:
    """
    Partition *frame* into individual chains.

    A chain ends at any line whose record name (cols 16) equals TER,
    irrespective of the remaining content (serial, residue ID, ).
    Consecutive TER lines are ignored; no empty chains are emitted.
    """
    chains, current = [], []

    for line in frame.splitlines():
        record = line[:6].strip()
        if record == "TER":               #  chain boundary
            if current:
                chains.append("\n".join(current))
                current = []
            continue                      # discard the TER itself
        if record == "ENDMDL":            # handled upstream
            continue
        current.append(line)

    if current:                           # flush final chain
        chains.append("\n".join(current))

    # Quality control  identical C count in every chain
    ca_counts = [
        sum(1 for l in c.splitlines()
            if l.startswith("ATOM") and l[12:16].strip() == "CA")
        for c in chains
    ]
    if len(set(ca_counts)) != 1:
        raise ValueError(
            f"Inconsistent C counts between chains: {ca_counts}"
        )

    return chains

def compute_ca_distances(chains: List[str]) -> np.ndarray:
    """Vectorised CC distances between *adjacent* chains."""
    coords_per_chain = []
    for chain in chains:
        coords = [
            (float(l[30:38]), float(l[38:46]), float(l[46:54]))
            for l in chain.splitlines()
            if l.startswith("ATOM") and l[12:16].strip() == "CA"
        ]
        coords_per_chain.append(np.asarray(coords, dtype=np.float64))

    n_residues = coords_per_chain[0].shape[0]
    if any(c.shape[0] != n_residues for c in coords_per_chain):
        raise ValueError("Chains do not contain the same number of residues.")

    # Trim termini
    s, e = BUFFER_RES, n_residues - BUFFER_RES
```

```
    coords_per_chain = [c[s:e] for c in coords_per_chain]

    # Pairwise distances between successive chains
    return np.concatenate([
        np.linalg.norm(coords_per_chain[i] - coords_per_chain[i + 1], axis=1)
        for i in range(len(coords_per_chain) - 1)
    ])

def main(pdb_path: str) -> None:
    frames = read_pdb_frames(pdb_path)
    if not frames:
        sys.exit("No MODEL/ENDMDL pairs found.")

    matrix = np.vstack([
        np.concatenate(([idx], compute_ca_distances(split_chains(frame))))
        for idx, frame in enumerate(frames, start=1)
    ])

    stem      = Path(pdb_path).stem                       # file name without .pdb
    directory = Path(pdb_path).parent                     # keep original directory
    out_path  = directory / f"{stem}_no_border_CA_all_distances.tab"
    np.savetxt(out_path, matrix, fmt="%.4f", delimiter="\t")

    n_frames, n_pairs = matrix.shape
    print(f"Saved {n_frames} frames  {n_pairs - 1} distances  {out_path.name}")

if __name__ == "__main__":
    if len(sys.argv) != 2:
        sys.exit("Usage:  python script.py  <input.pdb>")
    main(sys.argv[1])
```
